# Cellular insights of beech leaf disease reveal abnormal ectopic cell division of symptomatic interveinal leaf areas

**DOI:** 10.1101/2023.06.22.546113

**Authors:** Paulo Vieira, Mihail R. Kantor, Andrew Jansen, Zafar A. Handoo, Jonathan D. Eisenback

## Abstract

The beech leaf disease nematode, *Litylenchus crenatae* subsp. *mccannii,* is recognized as a newly emergent nematode species that causes beech leaf disease (BLD) in beech trees (*Fagus* spp.) in North America. Changes of leaf morphology induced by BLD can provoke dramatic effects into the leaf architecture and consequently to tree performance and development. The initial symptoms of BLD appear as dark green interveinal banding patterns of the leaf. Despite the fast progression of this disease, the cellular mechanisms leading to the formation of such type of aberrant leaf phenotype remains totally unknown. To understand the cellular basis of BLD, we employed several microscopy approaches to provide an exhaustive characterization of nematode-infected buds and leaves. Histological sections revealed a dramatic cell change composition of these nematode-infected tissues. Diseased bud scale cells were typically hypertrophied and showed a high variability of size. Moreover, while altered cell division had no influence on leaf organogenesis, induction of cell proliferation on young leaf primordia led to a dramatic change in cell layer architecture. Hyperplasia and hypertrophy of the different leaf cell layers, coupled with an abnormal proliferation of chloroplasts specially in the spongy mesophyll cells, resulted in the typical interveinal leaf banding. These discrepancies in leaf cell structure were depicted by an abnormal rate of cellular division of the leaf interveinal areas infected by the nematode, promoting significant increase of cell size and leaf thickness. The formation of symptomatic BLD leaves is therefore orchestrated by distinct cellular processes, to enhance the value of these feeding sites and to improve their nutrition status to the nematode. These results revealed a high specialized mode of parasitism of *L. crenatae* subsp. *mccannii*.

## Introduction

Beech leaf disease (BLD) is an emerging forest disease affecting beech trees (*Fagus* spp.) in North America. The first symptoms of BLD on American beech (*F. grandifolia* Ehrh) were first reported in Lake County Metroparks, located in the state of Ohio in 2012 (Ewing et al. 2019). The disease has rapidly spread to 11 other states throughout the eastern forest areas of the United States, as well as in Ontario, Canada (Ewing et al. 2019; Carta et al. 2020; Marra and LaMondia, 2020; Reed et al. 2020; Kantor et al. 2022a; Vieira et al. 2023). *Fagus grandifolia* is an important native deciduous hardwood forest species in North America. Together with sugar maple (*Acer saccharum* Marshall), and yellow birch (*Betula alleghaniensis* Britton), this group of trees are a dominant ecosystem of the northeastern United States and southern Canada hardwood forests (Bédard et al. 2022). As foundational tree species, beech trees play important roles in forest ecosystems, such as nutrient cycling, erosion control carbon storage and sequestration (Stephanson and Coe, 2017; Bayat et al. 2012; Schneider et al. 2015). Furthermore, its hardwood habitat characteristic (i.e., cavities and canopies) are crucial to provide nesting sites and shelter, and its nuts constitute a food source essential to the survival of several wild-life vertebrates (Leak et al. 2014). In addition to *F. grandifolia,* BLD also affects European beech, (*Fagus sylvatica* L.), Chinese beech (*Fagus engleriana* Seemen ex Diels) and Oriental beech (*Fagus orientalis* Lipsky) (Burke et al. 2020).

Beech leaf disease progression has accelerated in recent years in North America and has the potential to drastically alter these forest ecosystems (Zhao et al. 2023). Although BLD has been reported so far only in North America, there is an international concern regarding its potential spread to other areas around the globe (EPPO, 2018). For example, European beech (*F. sylvatica* L.) is an important forest tree and large natural resource in Western and Central Europe, broadly used for wood production (Ellenberg, 1988; Bolte et al. 2007). A critical monitorization plan has been established regarding the plant health status of *Fagus* spp. and possible risk of BLD introduction and spread within native beech forests in Europe (project FAGUSTAT).

Leaves are the primary organs for photosynthesis, and therefore, they play a vital role for plant growth and development (Kalve et al. 2014). The leaf is composed of a diverse set of cell types organized in three main tissues, i.e., mesophyll, vasculature, and epidermis (Tsukaya, 2013). Each group of cells has complex metabolic functions related to photosynthesis, gas exchange, and transportation of water and nutrients for the leaf, and ultimately for the full plant. Beech leaf disease induces dramatic changes on leaf morphology (Ewing et al. 2019; Burke et al. 2020), largely impacting the normal physiology of the leaf (McIntire, 2023). The initial symptoms of BLD appear as dark green interveinal banding patterns of the leaves, which are the hallmark of this disease. The leaf banding patterns are manifested at bud break and do not progress through, implying that the major events leading to these unique morphological changes occur through leaf development within the bud (Ewing et al. 2019; Fearer et al. 2022). Under highly severe levels of infection, leaves become leathery and crinkled (Ewing et al. 2019). An indirect example of the physiological changes associated with BLD is the noteworthy reduction of root ectomycorrhizal fungal colonization associated with severe diseased BLD trees (Bashian-Victoroff et al. 2023). As the disease progresses in the following years, there is a significant reduction in bud/leaf survival leading to a substantial decrease in the tree canopy, and eventually tree mortality (Ewing et al. 2019; Martin and Volk, 2021; Reed et al. 2022).

The beech leaf disease nematode, *Litylenchus crenatae* subsp. *mccannii* (Lcm, Family Anguinidae), is currently recognized as the major causal agent of BLD (Carta et al. 2020). Experimental laboratory inoculations of Lcm in buds of *F. grandifolia* resulted in BLD symptomatic leaves (Carta et al. 2020). Due to the negative consequences of this disease, this nematode is currently considered one of the top 10 most important plant-parasitic nematodes (PPNs) in the US (Kantor et al. 2022b). The origin of Lcm is still under investigation. Members of this genus have been found only in Japan and New Zealand, highlighting the lack of knowledge we still have about this genus. *Litylenchus crenatae* subsp. *crenatae* was first described from Japan (Kanzaki et al. 2019), associated with sporadic symptomatic leaves of Japanese beech (*F. crenata* Blume). So far, there is no report of BLD in Japan. The other species reported for this genus is *L. coprosma* found in association with small chlorotic patches of leaves of *Coprosma repens* A. Rich. (Zhao et al. 2011) and *C. robusta* Raoul-Karamu (Xu et al. 2017) in New Zealand. Although Lcm has been widely associated with symptomatic beech leaves in North America, the disease complex is still poorly understood, and the occurrence of other microorganisms potentially involved in this distinctive disease are still under debate (Burke et al. 2020; Ewing et al. 2021).

The family Anguinidae comprises both mycophagous and PPNs (Siddiqi, 2000). The latter are obligate specialized parasites of higher plants, mosses, and seaweeds, on which they frequently induce swellings or galls in different organs of the host (Subbotin and Riley, 2012). Several species are considered quarantine pests due to their economically damage as parasites of food [e.g., barley (*Hordeum vulare* L.), garlic (*Allium sativa* L.), onion (*Allium sepa* L.), potato (*Solanum tuberosum* L.), rice (*Oryzae sativa* L.), wheat (*Triticum aestivum* L.)] and ornamental crops in different areas of the globe (Siddiqi, 2000; Subbotin and Riley, 2012). In addition, few other species are known as important vectors of bacterial strains (e.g., *Rathayibacter toxicus*), which produce corynetoxins leading to annual ryegrass toxicity and poisoning of livestock (Murray et al. 2017). As migratory nematodes, they can migrate on the surface of the host tissues, and therefore, are often found as parasites of the aerial parts of different host plants (e.g., leaves, stems, inflorescences, seeds), and less frequently, in roots (Subbotin and Riley, 2012). While some species have a broad range of hosts, others have narrow host specificity (Siddique, 2000). Nevertheless, the resilient dialogue of these nematodes with the different host plant tissues suggests that they have a specialized ability in the manipulation of the host cellular machinery, which often results in the induction of cell hyperplasia and hypertrophy of the nematode feeding tissues (Subbotin and Riley, 2012; Palomares-Rius et al. 2017).

A decade after the first detection of BLD, there are still many gaps in our knowledge regarding this new forest disease. The emergence of this disease induced by a previously unknown nematode species brings many challenges due to the lack of information about its origin, spread, pathogenicity, mode of action, and other key components. An understanding of the cell biology underlying the development of BLD is crucial to gain an overall context of the cellular mechanism(s) leading to such peculiar leaf disease. In this study, we captured the sequential, temporal cellular events associated with the development of BLD, and simultaneously attempted to postulate an in-depth, mechanistic overview of this disease. Our assessment via light and scanning electron microscopy revealed extensive cellular changes of nematode-infected buds and leaf tissues. These changes were associated with a fine modulation of the host cellular mechanism(s) to alter the bud and leaf morphology to enhance the value of these feeding sites and to improve their nutrition status to the nematode. Our approach not only provided a good insight into the development of the nematode feeding site, but also revealed several critical time periods for future molecular and physiological studies of BLD ontogeny.

## Results

### 1. Beech buds can harbor a highly variable number of nematodes

Beech leaf disease begins within the bud because the leaves are already symptomatic when they break through the bud in early spring (Reed et al. 2020; Fearer et al. 2022). Therefore, we quantified the nematode population dynamics within the buds at the beginning of the autumn, when senescent leaves fall, and the buds become the only leaf related structure until the following season. The number of nematodes per bud was counted to determine the variation among trees in a BLD naturally infected forest stand. Sixty buds were collected from randomly distributed BLD symptomatic beech trees, and their dimensions and weights were measured. As individual scales were dissected from the buds, nematodes were found between the inner scales, while absent in buds (n = 25) collected from non-symptomatic trees. Dissections of buds revealed a large range of nematodes (n = 53±13 to 18,033±491) at this time of the year. All nematode developmental stages were found within the buds, including eggs at various stages of embryonic development, juveniles and adults. Surprisingly, adult males were found in less than 20% of the buds. The absence of males, associated with large numbers of eggs, suggested that this species can reproduce asexually. Detailed morphological analyses of adult females found within the buds revealed two female morpho-types: 1) amphimictic females present in < 20% of the buds; and 2) putative asexual females present in all buds.

### 2. Bud scale morphology is greatly affected by beech leaf disease

Beech buds are composed from a network of organized, overlapping bud scales, which encase and protect the shoot apex and the developing leaf primordial tissues from the environment. These inner tissues are ultimately responsible for the formation of new leaves. Comparison of the outer scales of non-and nematode-infected buds did not reveal any sign of infection or any apparent morphological changes; however, one of the first apparent symptoms of the disease is the pronounced morphological alterations of the inner bud scales that is associated with the presence of nematodes (Fig 1A). These morphological changes were divided into five distinct phenotypical groups based on the severity and extent of the bud scale symptoms (Fig 1A). In slightly symptomatic bud scales (groups 1 and 2) the symptoms of infection were localized in restricted areas on the lower part of the scale (Fig 1A). In these cases, nematodes congregated preferentially at the base of the scale corresponding to the growth zone of the bud scale. As the number of nematodes increased within the bud, the affected surface area progressively increased. In buds containing a higher number of nematodes (groups 3 and 4), the inner bud scales were highly distorted and wrinkled. In extreme cases, dark necrotic-like symptoms were observed in a large percentage of the bud scale. As noted above, the total number of nematodes present in each individual bud was highly variable, and the magnitude of the morphological changes was strongly correlated with the load of nematodes within each bud (Fig 1B). In addition, buds containing large numbers of nematodes (group 4) often revealed significant damage of the inner bud tissues, as shown in Fig 1C-1D.

**Fig 1.**
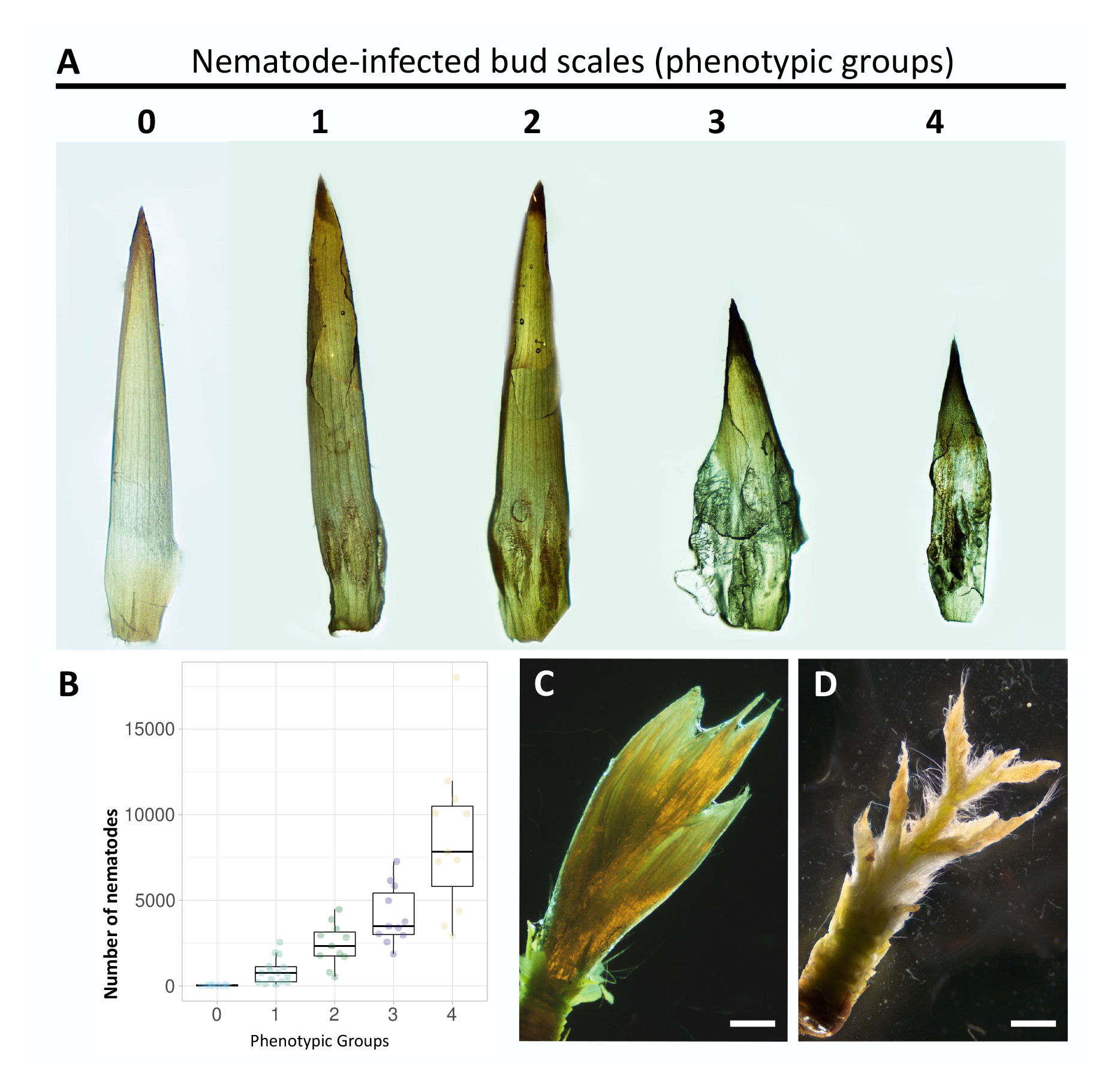
Bud scale morphology is dramatically affected by the presence of *Litylenchus crenatae* subsp. *mccannii*. **(A)** Morphological changes of bud scales infected with nematodes in early fall. A scale of 5 groups were established based on bud scale phenotype: (0 = no damage to 4 = significant damage). **(B)** Number of nematodes from 60 buds randomly collected from symptomatic beech leaf diseased trees distributed in a natural forest stand in Virginia (US). Buds were distributed within each phenotypic group (0 to 4). **(C-D)** Representative photomicrographs of the inner tissues of (C) non-infected and (D) highly infected buds. Scale bars: 2 mm.

Remarkably, these nematode-infected bud scales were characterized with irregular swellings (Fig 2D-F) compared to non-infected buds (Fig 2A-C). These swellings corresponded to hypertrophied cells and were depicted by their irregular cylindrical-like shape (Fig 2E-2F), when compared to the rectangular shape of normal cells of the bud scale (Fig 2B-2C). The size of these abnormal cells was highly variable within the same bud scale, or among different scales from the same bud. Large number of active nematodes were found between the compacted bud scales (Fig 2G). Individual nematodes were often observed with stylets retracted or protruded, apparently browsing on these enlarged scale cells that contained visible granular cytoplasm and cytoplasmic strands; potentially acquiring nutrients from these cells (Fig 2H). A significant number of eggs could be also found between the bud scales, suggesting that nematodes were actively reproducing at this time of the year (Fig 2I).

**Fig 2.**
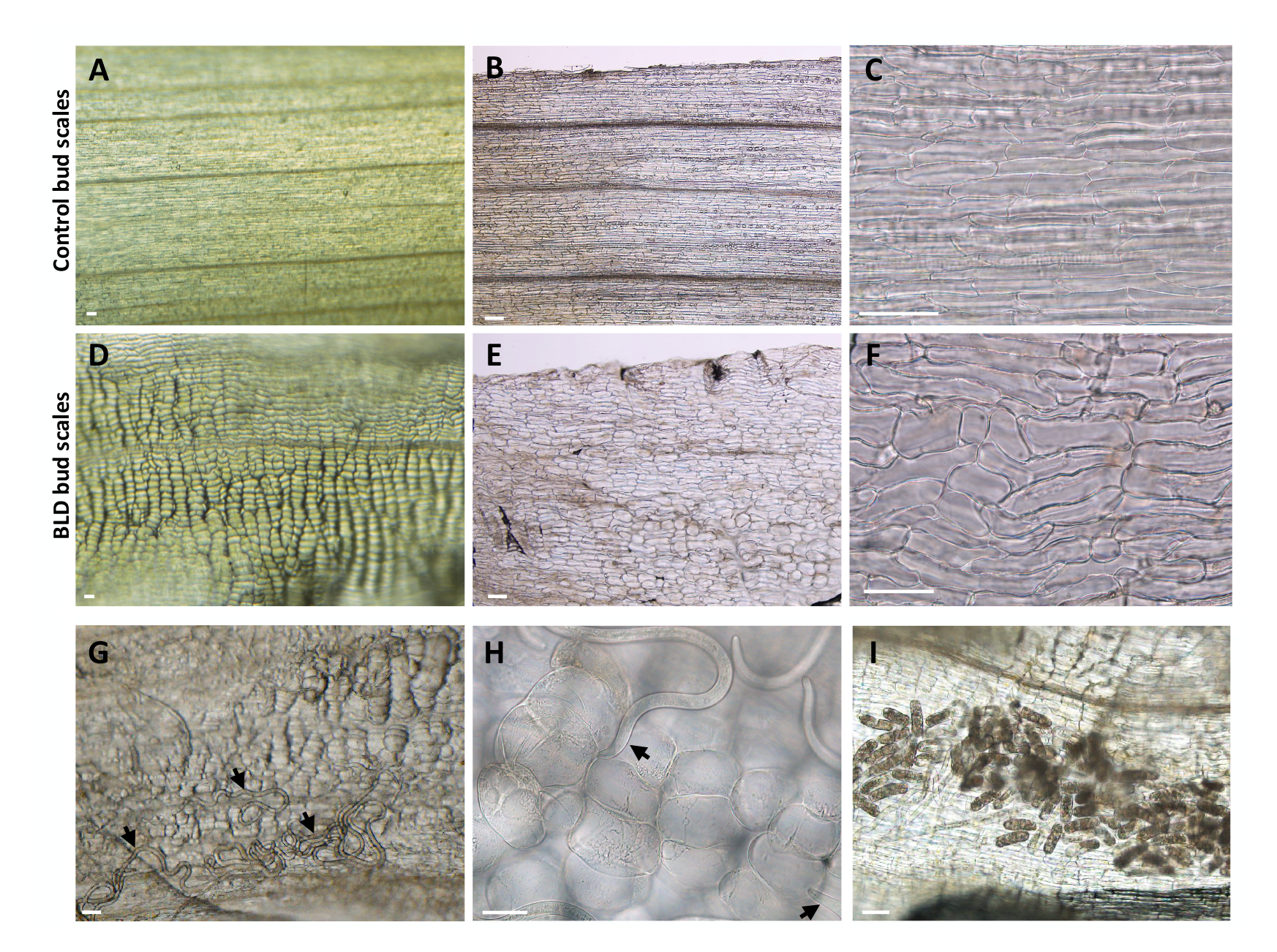
Nematode-infected (*Litylenchus crenatae* subsp. *mccannii*) bud scale cells showed variable levels of hypertrophy. **(A-C)** Photomicrographs of control bud scales showing flat, rectangular-shaped cells. **(D-F)** Representative infected bud scale cells. Infected cells presented several levels of hypertrophy and a more disorganized distribution of the cells (F). **(G)** Different nematode stages (arrows) associated with abnormally large scale cells. **(H)** Example of nematodes (arrows) probing large scale cells. **(I)** Large numbers of eggs were often found in nematode infected scales. Scale bars = 50 µm.

Bud scale morphology was examined at the cellular level using histological cross-sections (Fig 3), validating our previous observations (Fig 2). Control bud scales have 4-6 layers of relatively small cells (Fig 3A). Diseased scales had a similar number of cell layers; nevertheless, cells adjacent to nematodes were typically hypertrophied and showed a high variability of size that was probably related to their stage of development during the nematode interaction (Fig 3B-F). In some areas, cells were largely misshapen and distorted (Fig 3D-3F). Cells adjacent to nematodes were the most often affected; however, to a less extent, inner cells also reveal some levels of hypertrophy. Measurements of these larger cells using the surface area of fresh (Fig 3G) and fixed scale cross-sections (Fig 3H) confirmed the variability and significant increase in cell size in comparison to non-affected cells. Hypertrophied cells had a prominent nucleus and nucleolus, suggesting high metabolic activity. Measurements of the nuclear surface revealed up to a 5- fold increase in these larger cells compared to non-symptomatic cells (Fig 3I). Altogether, the occurrence of large nuclei associated with a generalized expansion of the affected bud scale cells suggested a possible link to the activation of the cell endoreduplication machinery.

**Fig 3.**
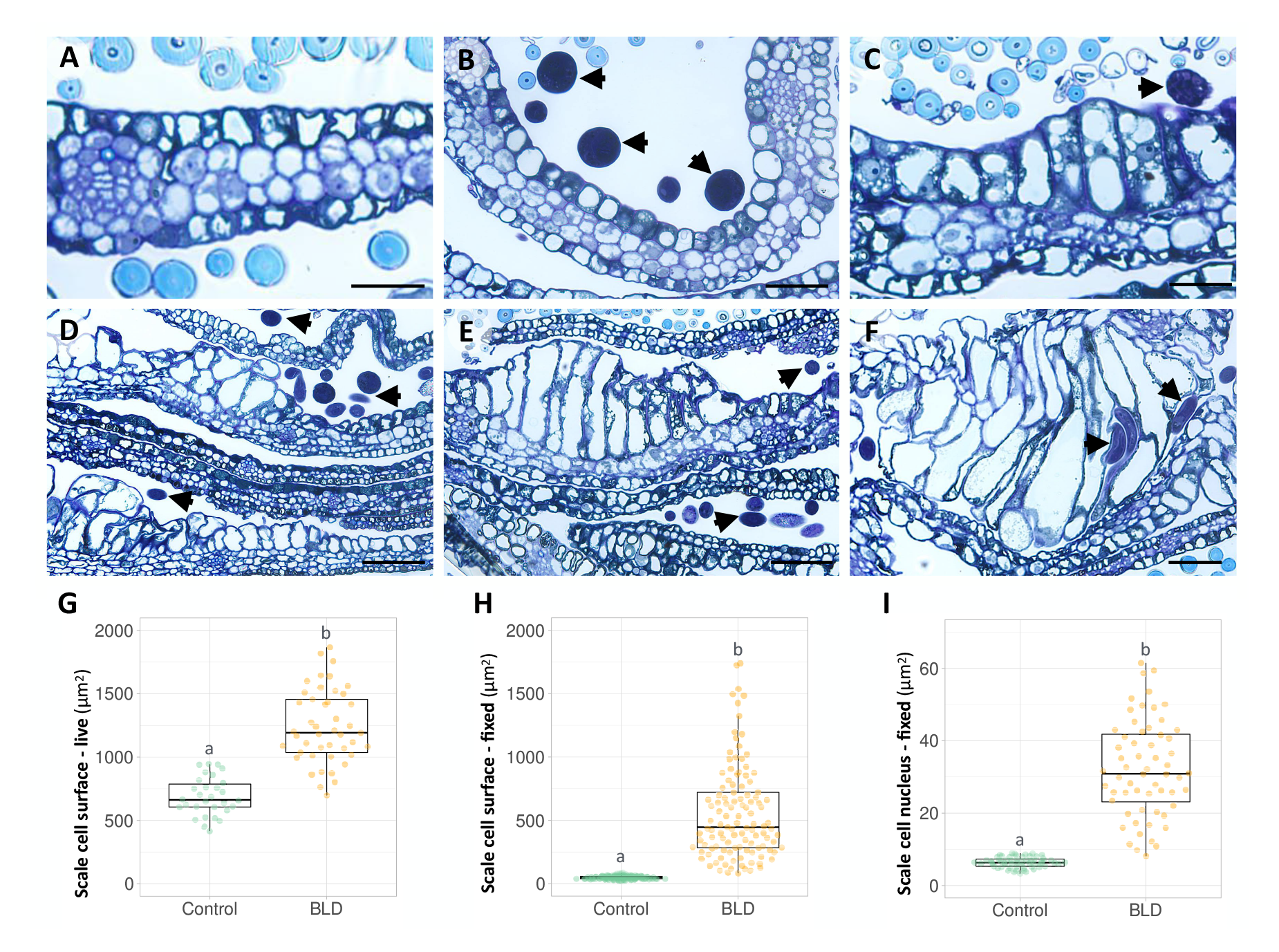
Histological analyses of non- and nematode-infected (*Litylenchus crenatae* subsp. *mccannii*) bud scales of beech (*Fagus grandifolia*). Bright-field photomicrographs of bud scale cross-sections stained with a mixture of toluidine blue and basic fuchsin. **(A)** Representative image of non-infected bud scale composed of four layers of cells. Spherical hyaline structures are sections through leaf hair. **(B-F)** Representative photomicrographs of nematode-infected bud scale cells showing different developmental stages of hypertrophy. Non-infected bud scale cells can also be seen adjacent to hypertrophied cells, as shown in figures C-F. Arrows point to nematode sections associated with hypertrophied bud scale cells. **(G)** Bud scale cell surface (µm^2^) was measured in control (n = 40) and nematode-infected (n = 60) whole fresh bud scales. **(H)** Bud scale cell surface (µm^2^) was measure in control (n = 115) and representative nematode-infected (n = 115) fixed and bud scale cross-sections. **(I)** Nuclear surface (µm^2^) of bud scale cell sections of control (n = 60) and nematode-infected (n = 60) buds. Box plots illustrate the means ± SEM. Different letters indicate statistical significance (*P* < 0.05). Scale bars: 25 µm (A-C); 50 µm (D-F).

### 3. Nematode-infected buds present altered cell division patterns in leaf primordial tissues

Some PPNs (including other members of the family Anguinidae) induce specialized nematode-feeding sites, which often involve a form of *de novo* organogenesis and requires the interaction with meristematic cells as a starting point (Palomares-Rius et al. 2017). In *F. grandifolia* the leaf primordia are initiated within the buds in late summer and expand and mature during the spring and early summer of the following year (Dengler et al. 1975). To this end, we investigated whether nematodes could interact with the inner parts of the bud, including the shoot apex and leaf primordial cells from histological sections of buds collected in early fall. Interestingly, different numbers of nematodes could be found associated with the shoot apex and surrounding the leaf primordial cells (Fig 4A-4B), suggesting that these cells can be a source of nutrients for the nematodes.

**Fig 4.**
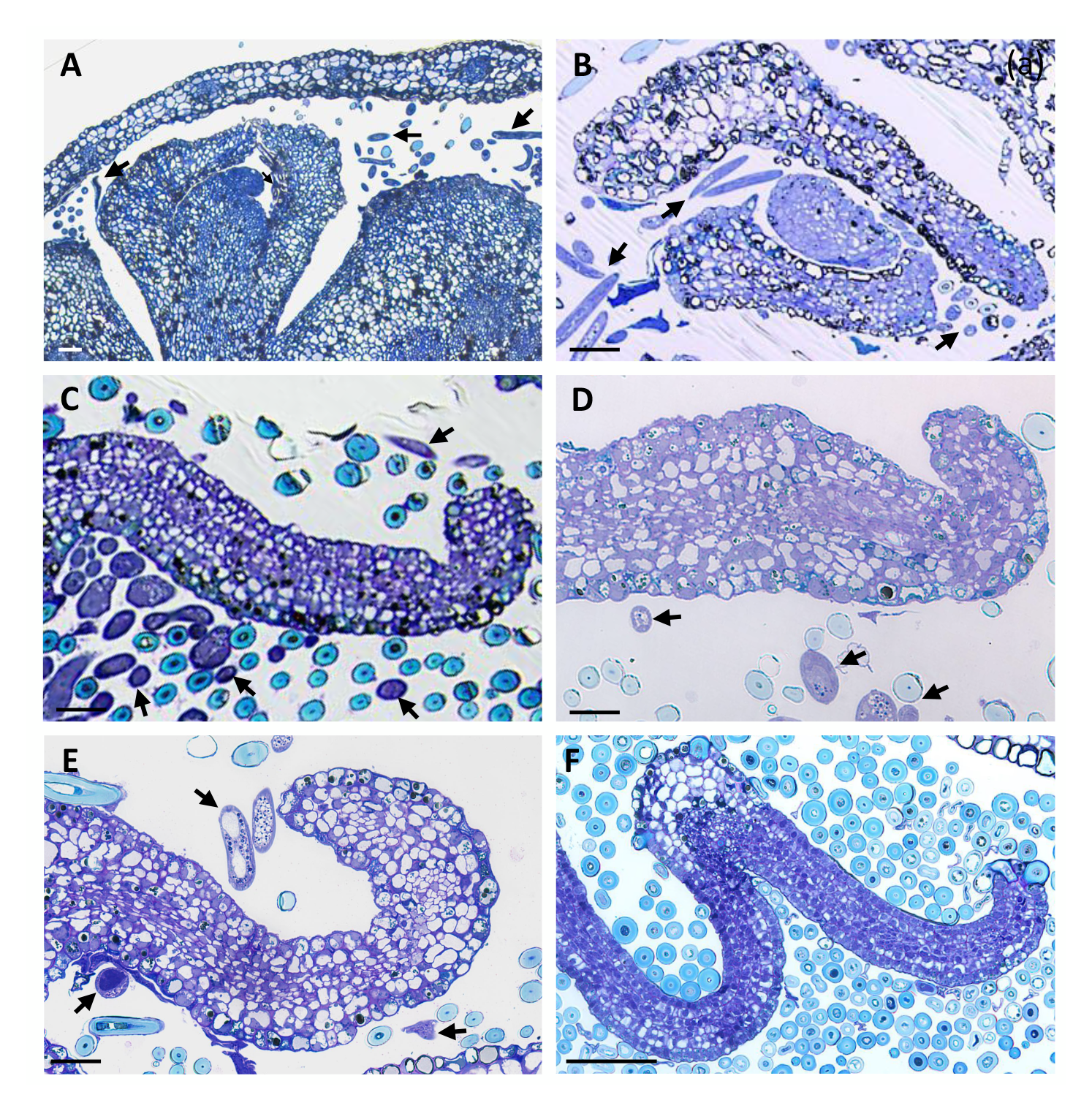
Histological analysis of shoot apex and leaf primordial tissues in non-and nematode-infected (*Litylenchus crenatae* subsp. *mccannii*) buds of beech (*Fagus grandifolia*). Bright-field photomicrographs of bud scale cross-sections stained with a mixture of toluidine blue and basic fuchsin are shown. **(A-B)** Beech leaf shoot apex and leaf primordial cells marking nematodes within these tissues (arrows). **(C-E)** Examples of leaf primordial developing in buds infected with nematodes (arrows). Note the presence of different number of disorganized and abnormal cell layers (n > 6) in each leaf primordia. **(F)** Cross-section of leaf primordia from a non-infected bud showing six distinct layers of highly organized cells. Scale bars: 50 µm.

Surprisingly, nematode-infected buds revealed a noticeable variability at the leaf primordial cell composition within and between individual buds. Instead of the typical six layers of highly organized cells reported for *F. grandifolia* (Dengler et al. 1975; Fig 4F), leaf primordium flanks associated with nematode-infected buds revealed an altered cellular patterning (Fig 4C-E). In these infected buds, the leaf primordium flanks could vary from six to more than ten cell layers showing different levels of disorganized arrangements. Cell size was also highly variable in leaf primordium associated with nematodes, resulting in thicker leaf tissue. These discrepancies in leaf primordial cell structure suggests that a differential rate of cellular division occurs within areas of the leaf in nematode-infected buds, leading to the formation of additional cell layers of the leaf in those areas.

### 4. Assessment of nematode and leaf development in dormant buds

Buds become dormant with the decrease of temperatures and photoperiod in the fall (Falusi and Calamassi, 1977). During dormancy there is a gradual metabolic and physiological arrestment of the apical meristem and the young developing leaves inside the bud until the following spring (Ruttink et al. 2007). On the other hand, the life cycle of PPNs is often constrained by the seasonality of resources coupled with unfavorable abiotic conditions (Siddiqi, 2000). To evaluate both nematode and leaf development during the colder months, buds were collected at two different time points in the winter (i.e., December and late February). In December, 20 buds were dissected to assess only nematode development in association with dormant buds. Nematodes could be often found aggregated as “nematode wools” between the bud scales, a typical behavior often seen in PPNs when expose to adverse abiotic conditions. Although buds can provide a favorable micro-habitat from hostile environment conditions, the transition of seasons was marked by a switch of the nematode dynamics in the colder months. These transitions probably reflect the biology of this species upon exposure to cold conditions, which included: 1) absence of viable eggs, due to the decline of adult females (i.e., only remnants of adult females were found); 2) buds were populated mainly by juvenile stages with characteristic dark intestines filled with more lipid droplets. The most predominant active stage found was immature females. Based on these observations, and similar to nematodes found in senescence beech leaves (Kanzaki et al. 2018), the immature females were the main resistant and winter survival stage of this species.

To examine nematode population dynamics associated with leaf development in dormant buds, an analysis was performed by dissecting a set of 50 buds collected at the end of the winter season (Fig 5). As previously reported, the scale morphology was used as a marker for nematode infection. At this time of the year, the large bud scale cells associated with the presence of nematodes were flattened and presented a brown discoloration that was potentially associated with the long-term effect of the cold conditions (Fig 5A). Inspection of young leaves of less than 10 mm in length established within these buds revealed distinctly patterned symptoms that resembled the typical interveinal dark banding observed in mature BLD leaves. A set of representative leaves with few interveinal symptomatic areas to a more exacerbated and widespread distribution of the interveinal-like symptoms is shown in Fig 5B. Next, the correlation of the different leaf phenotypes and the load of nematodes within each bud was investigated.

**Fig 5.**
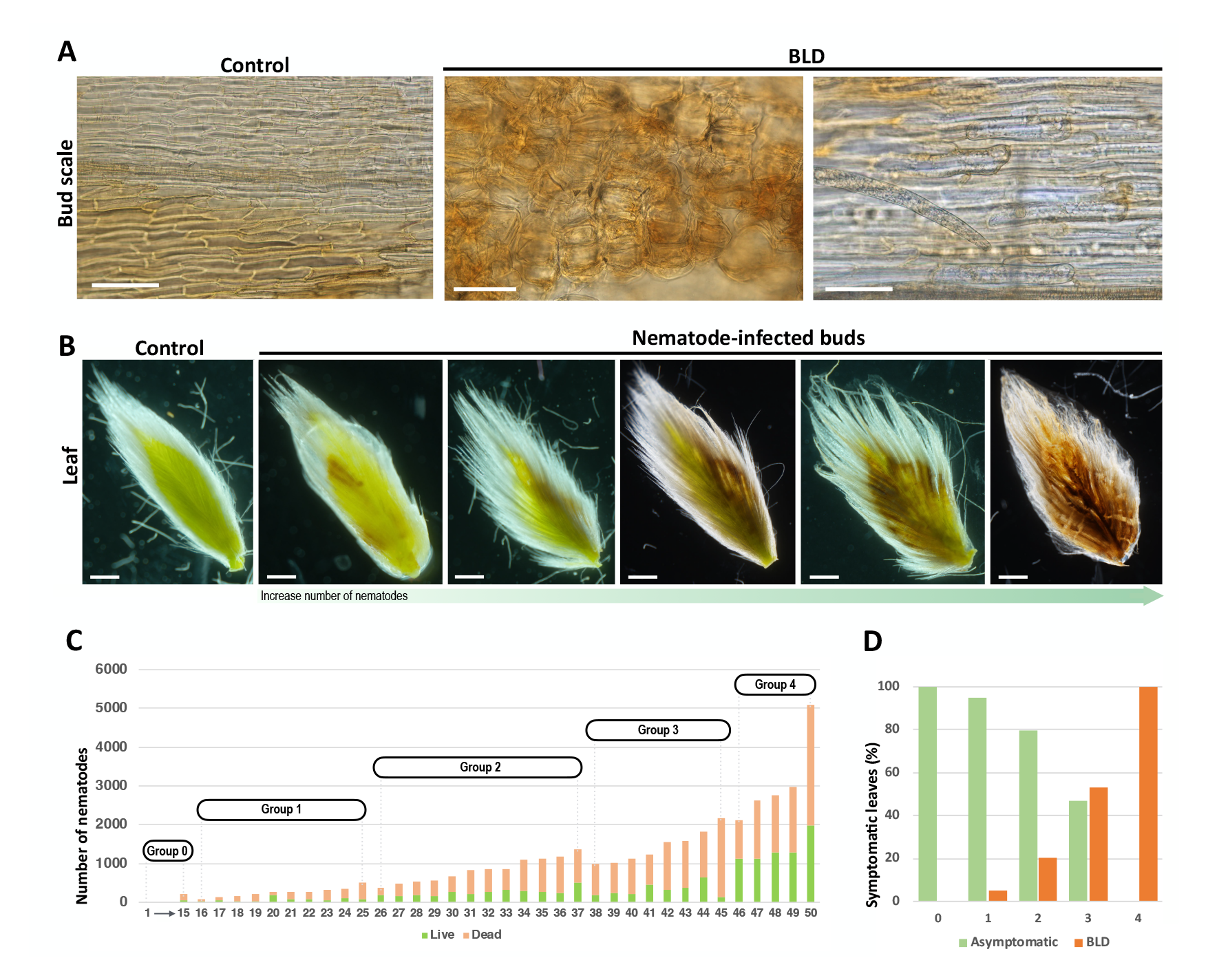
Nematode (*Litylenchus crenatae* subsp. *mccannii*) and leaf primordia development in dormant buds of beech (*Fagus grandifolia*). Fifty buds were randomly collected from symptomatic beech leaf diseased (BLD) trees in late February. **(A)** Representative bud scale cells of control and nematode-infected buds. Cell hypertrophy is associated with the presence of nematodes. **(B)** Representative leaves collected from control and nematode-infected buds. Buds infected with a high population of nematodes caused highly symptomatic and damaged leaves. **(C)** Number of nematodes quantified for 50 random buds. Both active and dead nematodes were quantified and represented by different colors in a bar graph. Buds were phenotyped according to the previously established groups (see Fig 1). **(D)** Percentage of asymptomatic and symptomatic BLD leaves associated with 50 dormant buds distributed by their phenotype. Scale bars: 50 µm (A); 1 mm (B).

At this time of the year, a large percentage of nematodes within the buds were dead, however, all nematodes were quantified (Fig 5C). The occurrence of these leaf morphological changes was strongly concurrent with the proportion of nematodes found within each bud and the severity of phenotypic effects varying between the buds. Buds with a smaller load of nematodes showed minor proportions of apparent symptomatic leaves (groups 1 and 2). In contrast, buds containing larger numbers of nematodes presented a high frequency of symptomatic leaves, ranging from 55% to 100% in groups 3 and 4, respectively (Fig 5D). It is noteworthy, leaves associated with buds from group 4 containing a higher number of nematodes, displayed severe necrotic-like symptoms, potentially inducing a detrimental effect, and compromising leaf development.

To follow the cellular changes associated with this altered leaf morphology, histological sections were made from control (Fig 6A) and nematode-infected buds (Fig 6B). Following our previous results, nematodes were found between the bud scales. In addition, nematodes were found between the interveinal areas bounded by leaf veins of the developing leaves (Fig 6B-F). The distribution and number of nematodes associated with the different interveinal areas varied not only between the interveinal areas of the same leaf, but also among the interveinal areas of different leaves developing within the same bud (Fig 6B). Strikingly, interveinal leaf areas with higher number of nematodes displayed newly formed cell layers, resulting in a significant increase of its thickness (Fig 6G-6H). In some cases, both interveinal and vein cells displayed a significant expansion when compared to non-infected leaf cells (Fig 6G-H), reinforcing the potential ability of nematodes of inducing larger cells. By contrast, and similar to control leaves (Fig 6C), non-infected interveinal areas, or areas with a reduced number of nematodes, were characterized by six distinct layers of orderly cells of relatively similar sizes. In line with the previous observations, the nuclear surface of these enlarged cells was also significantly increased when compared with the nuclei of non-infected cells. This type of ectopic cell proliferation in different interveinal portions of the leaf can be related to the typical BLD interveinal banding observed on mature leaves.

**Fig 6.**
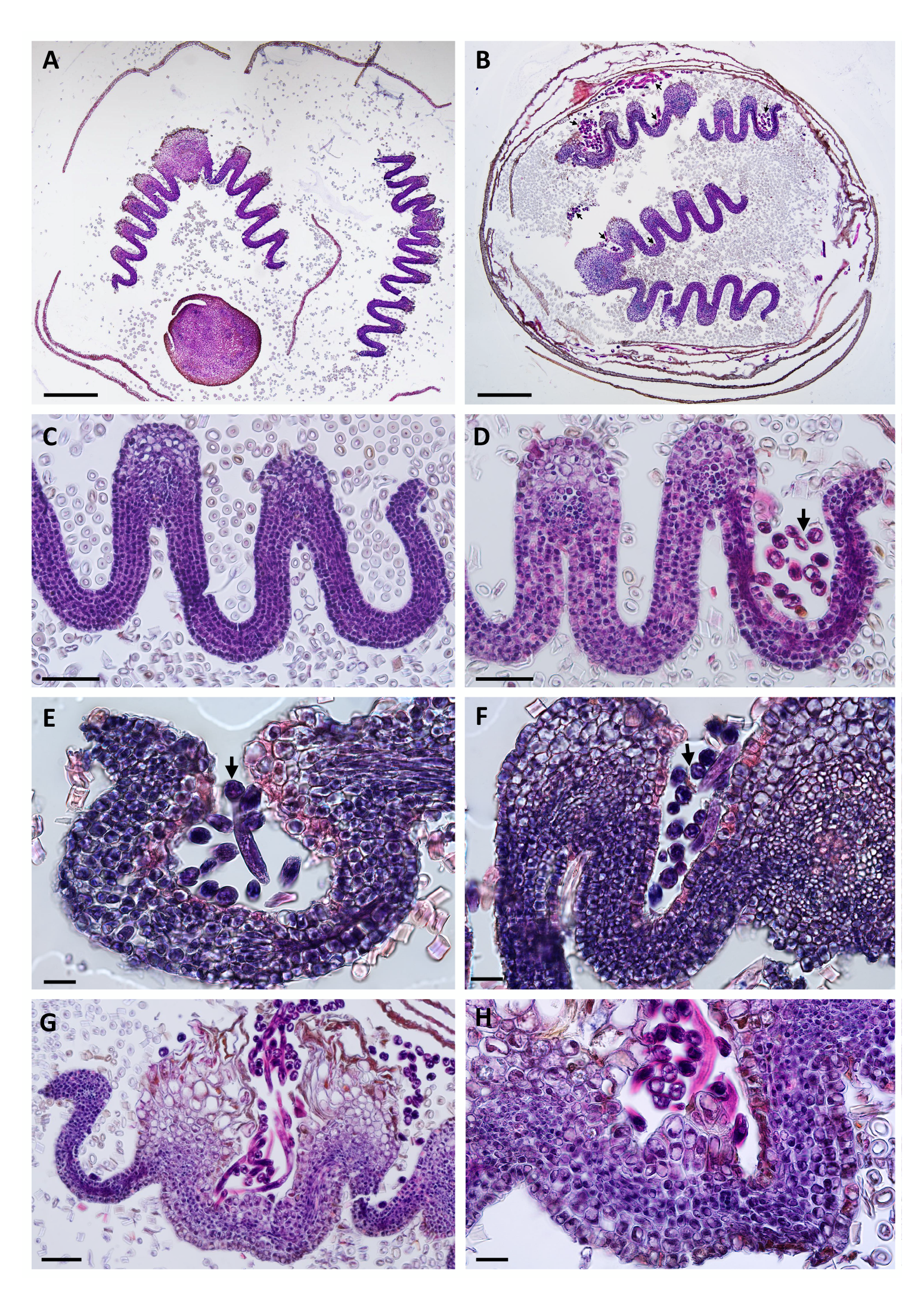
Histological sections of dormant beech (*Fagus grandifolia*) buds infected with *Litylenchus crenatae* subsp. *mccannii*. Control and nematode-infected buds were collected at the end of February. Bright-field photomicrographs of bud scale sections stained with a mixture of hematoxylin and eosin. **(A)** Cross-section of a control dormant bud showing two developing leaves. Circular hyaline structures are sections of leaf hairs. **(B)** Cross-section of a dormant nematode-infected bud. Different number of nematodes (arrows) were found between the bud scales, as well as in the interveinal areas of the leaf. **(C)** Higher magnification of the section of a control leaf showing the typical six layers of cells. A large number of leaf hairs are often present in these leaf sections. **(D-F)** Higher magnification of the sections of interveinal leaf areas with nematodes (arrows). **(G-H)** Nematode-infected interveinal beech leaf area showing morphological changes of augmented cell division. Scale bars: 200 µm (A-B); 50 µm (C, D, G); 20 µm (E, F, H).

The earlier observations demonstrated that the abnormalities in the external leaf morphology begin at earlier times of leaf development (i.e., in leaves that are a few mm long). This prompted us to evaluate in more detail the cell anatomy of individual leaves associated with these nematode-infected dormant buds (Fig 7). As previously reported for *F. grandifolia* (Dengler et al. 1975), leaves dissected from non-infected buds comprised four layers of mesophyll cells flanked by single cell layers on both adaxial and abaxial epidermis as shown in Fig 7A. In contrast, sections of nematode-infected interveinal leaf areas displayed major changes in their cellular composition as represented in Fig 7B-D. In this leaf, while some interveinal areas displayed only slightly changes on the number of cells and cell layers, others displayed a dramatic increase of cell proliferation and an overall disorganized cell pattern, suggesting the occurrence of differential and localized cell division rates in different parts of the leaf. This upsurge of cell and lack of cell order is apparent when we traced the cell layer outlines on a series of interveinal cross-sections of the same leaf (Fig 7B’-7C’) in comparison to control (Fig 7A’). We also noted that this ectopic cell proliferation could occur along different veins of the leaf, resulting in a dramatic change of their overall cell architecture (Fig 7E). Whereas this increase of cell layers was restricted and punctually localized in some interveinal areas of the leaf, in other nematode-infected buds, the cell layers surge was more intensified covering larger interveinal leaf areas (Fig 7F). Hence, the extent of induced cell layers varied in different areas of the leaf, as well as the number of apparent smaller cells within these areas.

**Fig 7.**
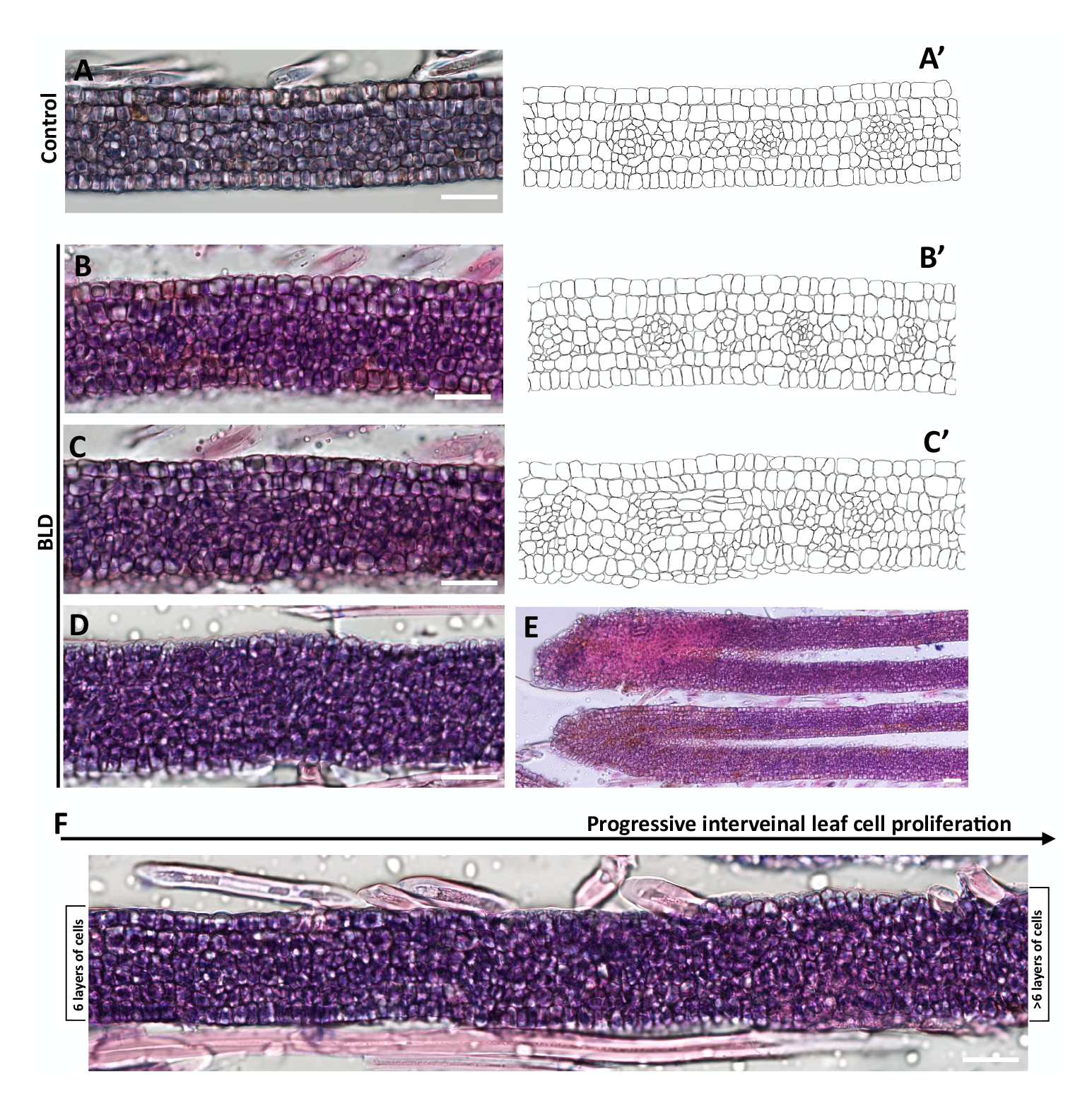
Sections of young developing leaves within nematode-infected (*Litylenchus crenatae* subsp. *mccannii*) beech buds (*Fagus grandifolia*) with altered interveinal leaf cell proliferation. Morphological cell rearrangements of interveinal leaf cross-sections. **(A)** Representative section of control leaf interveinal areas with the typical six layers of cells. **(B-D)** Three interveinal areas of the same leaf infected by nematodes showing a variable and prominent increase in the number of cells layers. Outline drawings of some representative interveinal areas highlighting this upsurge in cell division associated with nematodes (B’-C’) compared to the control (A’). **(E)** Veinal areas of the same nematode-infected leaf displayed different levels of ectopic cell proliferation. **(F)** Differential cell proliferation within the same interveinal leaf area dissected from a nematode-infected bud. Scale bars: 20 µm.

### 5. Penetration of emergent leaves by nematodes in the spring

Foliar nematodes, including species within the Anguinidae, are capable of migrating externally in films of water on petioles, stems, and foliar surfaces (Siddiqi, 2000; Subbotin and Riley, 2012). As *F. grandifolia* leaves breakout through the buds in the spring, leaf cells initiate their expansion and maturation (Dengler and MacKay, 1975; Dengler et al. 1975). Therefore, we questioned the nematode transition mechanism from the buds into new fully exposed leaves. Because of the variation in leaf size at this time of the year (Fig 8A), dozens of symptomatic leaves of different sizes with BLD symptoms were stained with acid fuchsin (Fig 8B). Notably, individual and groups of nematodes were found migrating and penetrating the abaxial surface of the leaves (Fig 8C). In some cases, several nematodes used the same point of entry into the leaf. Nematodes were often associated with the development of minute necrotic areas at the leaf surface. Despite nematodes being found in the surface of maturing leaves, the total number of nematodes recovered from emergent symptomatic leaves at this time of the year was very restricted (i.e., varying from 9 to 216 nematodes detected per leaf; n = 30). These results were consistent with previous surveys reporting a reduced number of active nematodes recovered in the spring from symptomatic BLD leaves collected in other natural infected areas (Reed et al. 2000).

**Fig 8.**
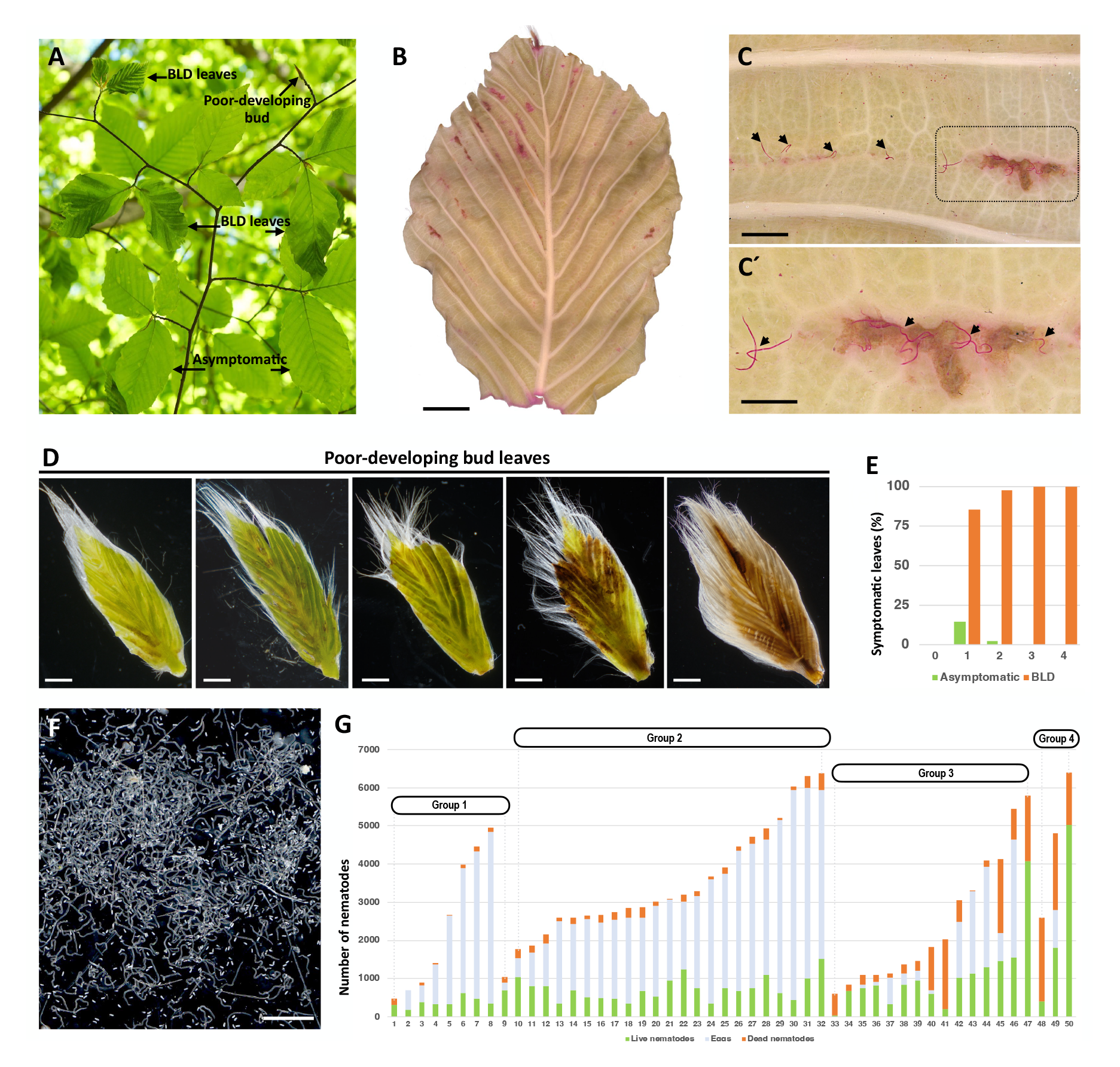
Nematode (*Litylenchus crenatae* subsp. *mccannii*) assessment of leaves and beech (*Fagus grandifolia*) buds in early spring. **(A)** Representative branch showing sets of asymptomatic and symptomatic beech leaf disease (BLD) leaves, and poorly developing buds. **(B-C)** Detection of nematodes in symptomatic BLD leaves using acid fuchsin staining (B). Nematodes (arrows) were found migrating on the abaxial side of the leaf and penetrating the leaf tissues (C-C’). **(D)** Representative symptomatic BLD leaves dissected from poorly developing buds. **(E)** Percentage of asymptomatic and symptomatic BLD leaves associated with 50 poor-developing buds rated according to the phenotype scale presented in Fig 1. **(F)** Active nematodes and eggs extracted from a single, poorly developing bud. **(G)** Numbers of nematodes extracted from 50 poorly developing buds. Buds were classified from 0 to 4 based on their bud scale phenotype (see Fig. 1). Scale bars: 5 mm (B); 1 mm (C, D, F); 500 µm (C’).

The incidence of poor-developing buds is a common phenomenon associated with symptomatic BLD trees (Ewing et al. 2019). To evaluate to what extent the occurrence of this type of buds’ correlates to the presence of nematodes, 50 buds were randomly collected from symptomatic BLD trees in early spring (Fig 8A). The inspection of individual buds confirmed the presence of nematodes in all the buds, revealing highly levels of infection. An extended assessment of the bud samples revealed visual symptoms for >95% of the internal leaves, comprising typical interveinal green banding and/or crinkled leaf morphology to highly necrotic-like and damage tissue (Fig 8D-8E). Remarkably, nematodes resumed their development and reproduction in most of the assessed buds, with females laying large number of eggs (Fig 8F). Moreover, the incidence of necrotic leaves correlated with a larger percentage of dead nematodes, as well as smaller number of eggs, in comparison to other less necrotic symptomatic BLD leaves (Fig 8G). Collectively, our findings underline the biotrophic relation of Lcm and the need of using living cells as source of nutrients as other PPNs.

The fact that nematodes migrate along the leaf surface prompted us to evaluate whether nematodes could also migrate in the twigs adjacent to highly infected buds. Over the 15 twigs assessed, which were composed of 5-6 buds each, live and dead nematodes were isolated from the surface of 10 (66.6%) twigs. Morphological and qPCR analyses confirmed the identification of Lcm. The migratory nature of these nematodes hinted towards the possible movement and infection of new emergent symptomatic BLD leaves, or other non-infected areas of the tree. More importantly, poor-developing buds can be considered as sources of high nematode inoculum.

### 6. Symptomatic BLD leaves display major cellular structural modifications

To gain insight into the cellular changes associated with mature symptomatic BLD leaves, we employed different approaches of microscopy to provide an exhaustive characterization of the leaf morphology. To this end, symptomatic leaves were collected at early autumn to capture all type of leaf symptoms within the same individual leaf, i.e., leaves containing both green and dark yellowish/brownish interveinal banding areas, as well as non-symptomatic areas. While in the spring we give priority to symptomatic crinkle leaves. First, we focused our analyses on the surface of the abaxial epidermis to evaluate any changes of morphology using light and scanning electron microscopy (SEM) since nematodes were seen migrating along these tissues. When examined under low magnification, control leaves were characterized by a flat lamina with a complex reticulate venation architecture (Fig 9A). However, interveinal green symptomatic leaf areas were characterized by moderate swellings on the abaxial side of the leaf, partially covering the interveinal pattern structure (Fig 9B). In acute symptomatic leaf areas (yellowish/brownish coloration) the interveinal venation architecture become totally indistinct with significant increases in swelling (Fig 9C). Similarly, crinkle leaves also displayed moderate swellings of the adaxial surface of the leaf (Fig 9D).

**Fig 9.**
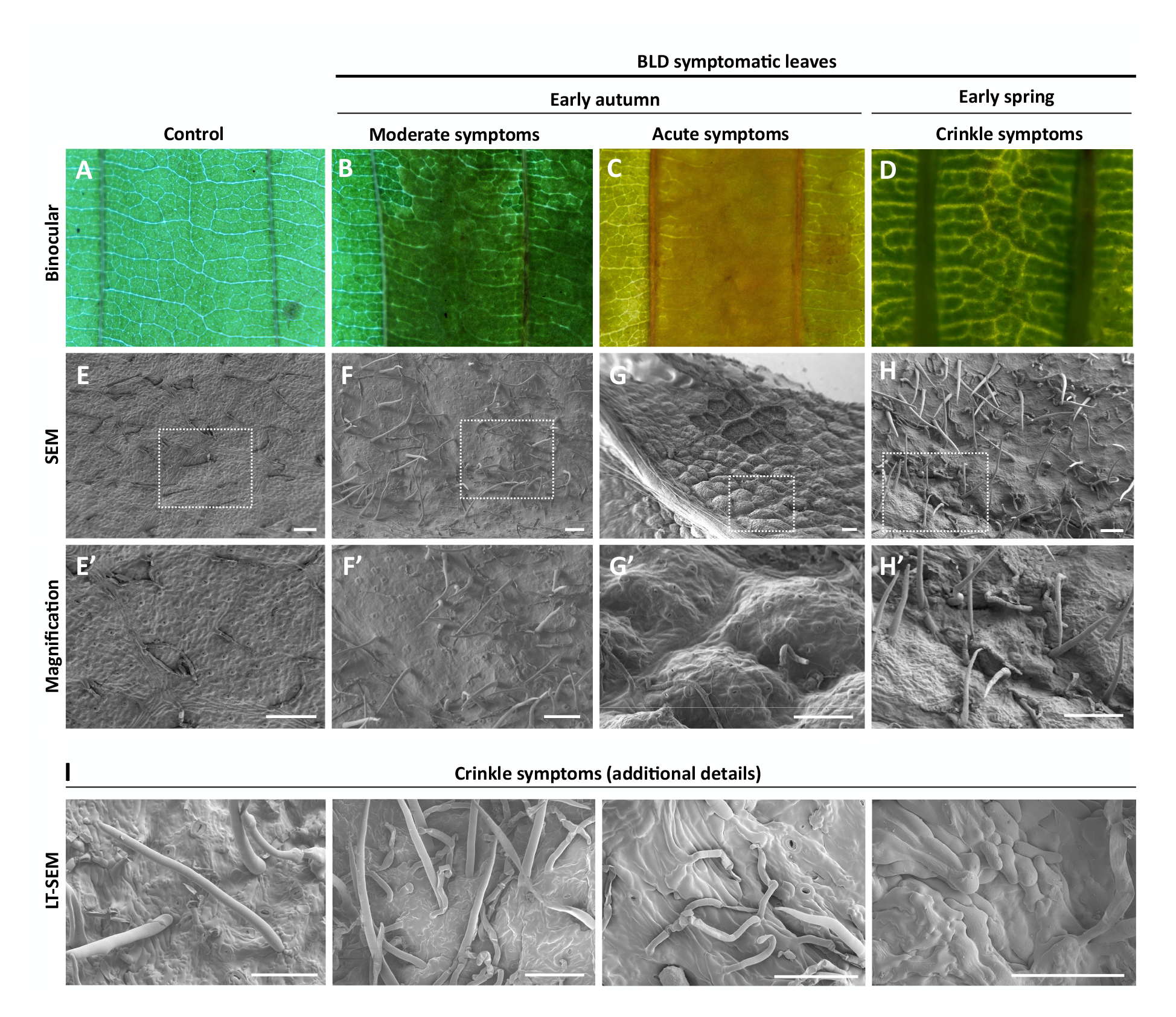
Structural characterization of abaxial leaf surface of beech (*Fagus grandiflora*). **(A-D)** Abaxial interveinal areas of control (A) and symptomatic beech leaf disease (BLD) leaves (B-D) showing different interveinal banding were observed with a stereoscope. **(E-H)** Scanning electron microscopy (SEM) photomicrographs of abaxial interveinal areas of control (A) and symptomatic BLD leaves (E-H). Note the progressively swollen tissues within the interveinal areas of symptomatic BLD leaves collected in early autumn (F-G). Crinkled leaves collected in early spring displayed significant ectopic cell proliferation. White squares within the photomicrographs are magnified in E’-H’. **(I)** Additional details of crinkled leaf morphology were observed using low-temperature SEM confirmed irregular ectopic cell division. Scale bars: 100 µm.

To allow observations at higher magnification, SEM was used to observe infected and non-infected leaves (Fig 9E-I). Control abaxial leaf surface displayed a flat lamina, with the presence of a significant number of stomata and the presence of trichomes (Fig 9E). In the dark green banded areas of the leaf, moderate swellings on the abaxial side were observed along the symptomatic interveinal areas (Fig 9F). BLD symptomatic areas were also noted to have a comparatively larger number of trichomes than non-infected leaves. Gradually, leaves with acute phenotypic symptoms revealed overgrowth pouch-like areas of the abaxial side of the leaf (Fig 9G). The outline of the abaxial side of the leaf revealed pronounced convexly curved protrusions. Often, patches of these pouch-like areas were not present in different individual processed leaves, probably as a consequence of the large internal pressure inside the leaf. Interestingly, crinkle leaves present dramatic ectopic cell division, resulting in overall aberrant leaf morphology (Fig 9H) and irregular cell surface composition with cells being stack in the top of each other (Fig 9I).

The occurrence of an abnormal increase of tissue in the abaxial side of the leaf prompted us to examination the cell morphology of the epidermal pavement cells of leaves collected in early autumn (Fig 10A-10B). Therefore, different areas abaxial epidermal cells of control (Fig 10C) and leaves with acute BLD symptoms were traced (Fig 10D). Whereas the epidermal pavement cells in uninfected leaves assumed the classical jigsaw puzzle pattern, those of nematode-infected leaves showed prominent cell-shape deformations as well as an apparent reinforcement in the cell wall. To establish whether these morphological changes correlate to the presence of nematodes, the chlorophyl was removed and the nematodes were stained with acid fuchsin. Remarkably, hundreds of nematodes could be observed within the inner layers of BLD symptomatic leaves (Fig 10F), while absent in control leaves (Fig 10E). Similar analyses were then performed in non-and symptomatic BLD areas within the same leaf (Fig 10G). Likewise, similar patterns of epidermal cell differentiation were observed, as well as a large number of nematodes associated with the symptomatic areas of the leaf.

**Fig 10.**
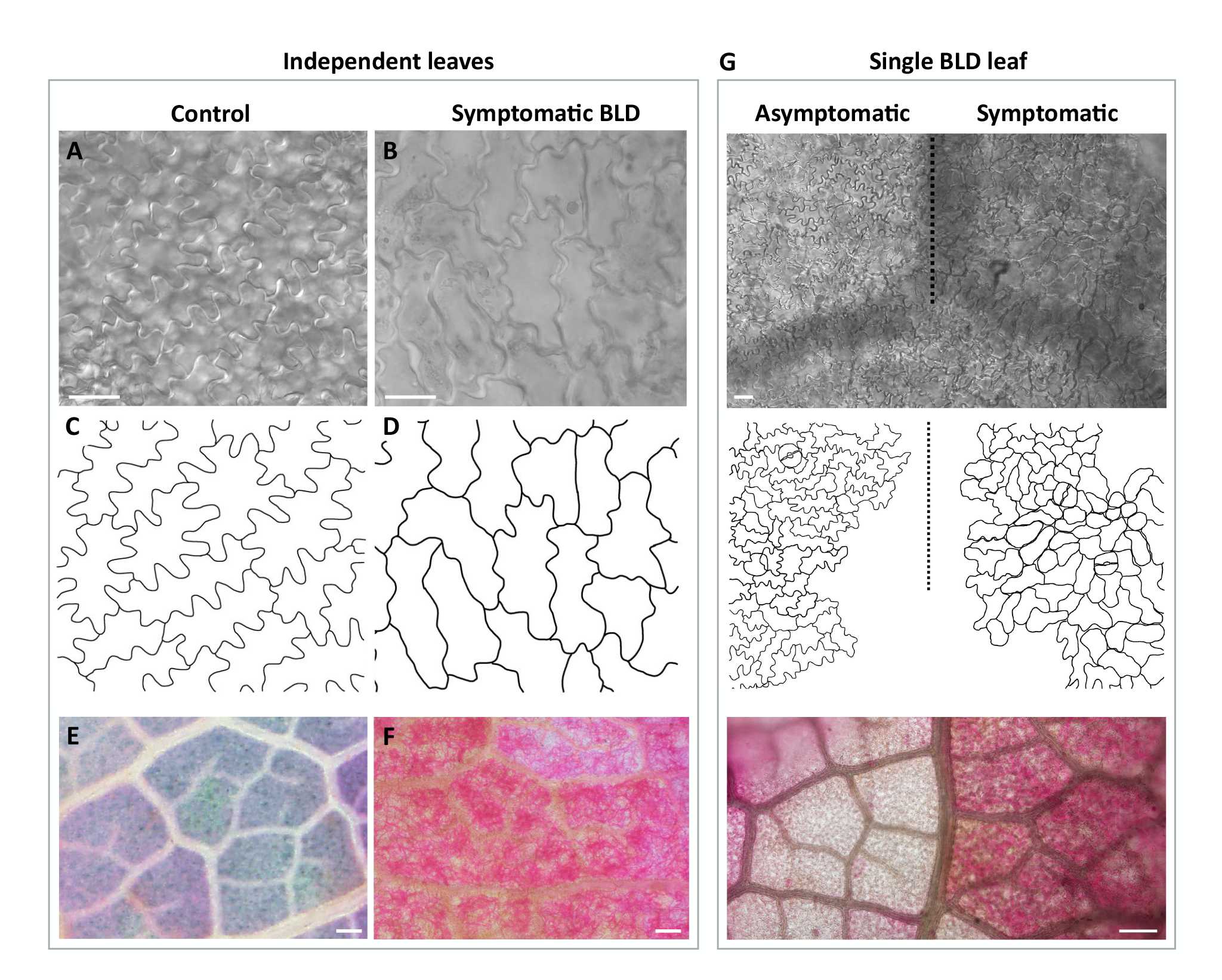
Abaxial epidermis pavement cells pattern of beech (*Fagus grandifolia*) leaves. Control and symptomatic beech leaf disease (BLD) leaves were collected in early autumn. **(A-B)** Abaxial epidermal pavement cells of control (A) and symptomatic BLD leaves. **(C-D)** Schematic representation of the pavement cells of control (C) and symptomatic BLD (D) leaves. **(E-F)** Acid fuchsin staining of control (E) and symptomatic BLD (F) leaves. Note the high number of nematodes stained within the leaf tissues (red). **(G)** The same type of analysis was performed using asymptomatic and symptomatic BLD areas of the same leaf. Scale bars: 20 µm (A, B, G upper panel); 200 µm (E, F, G lower panel).

### 7. Changes of symptomatic BLD leaf coloration is associated with nematode activity

The progression of BLD symptoms is typically characterized by gradual changes of the interveinal leaf coloration. While in the spring the BLD interveinal areas can present distinct shades of green, by the end of the summer some of these interveinal areas will turn yellow to brown-like necrotic areas (Fig 11A- C). We performed sections of fresh leaves containing both interveinal dark green and yellowish dark symptoms to establish the potential causal variation of BLD coloration. To this end, the spongy mesophyll was gently striped off in both symptomatic and un-infected leaves as shown in Fig 11D-F.

**Fig 11.**
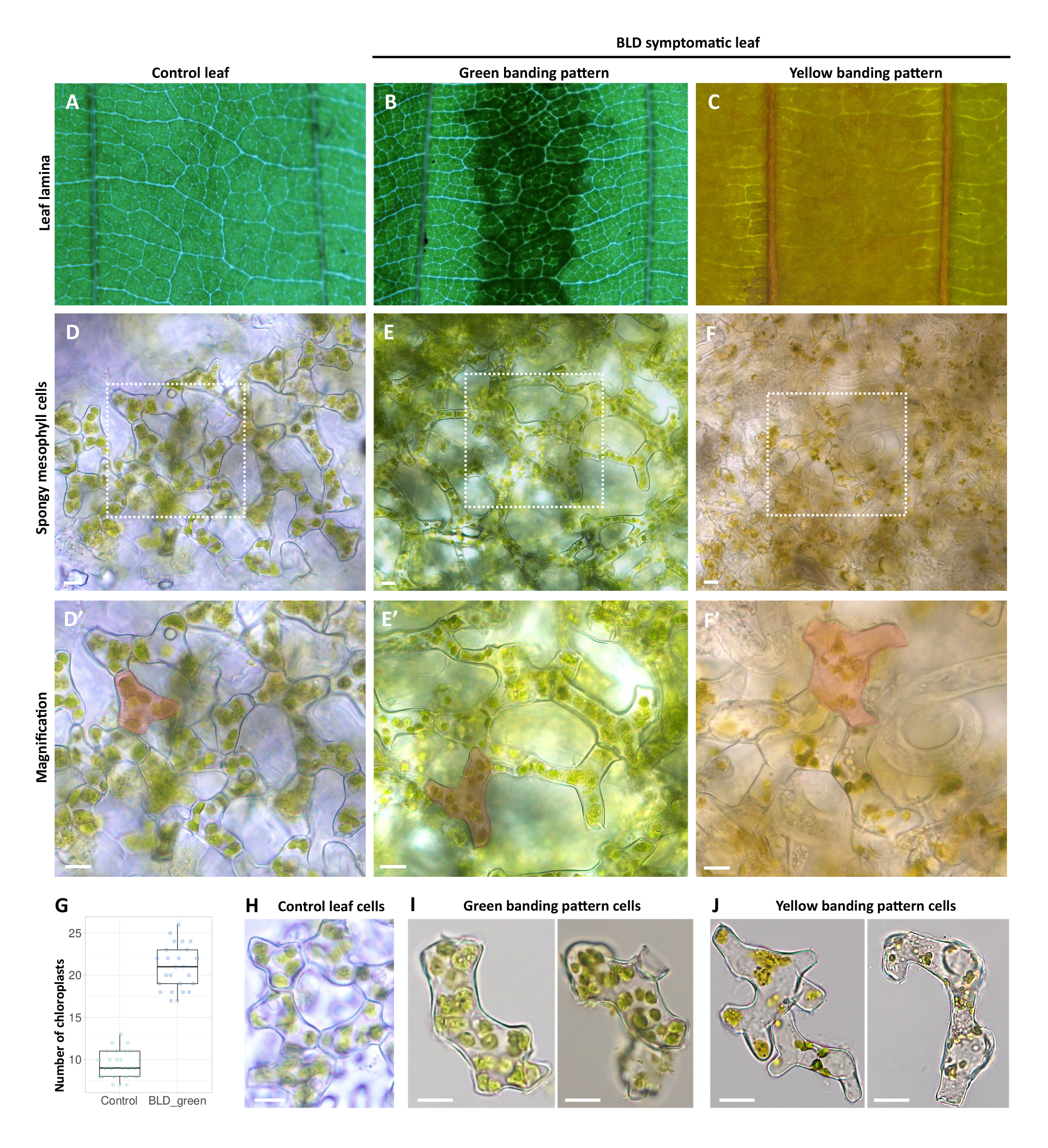
Life cell imaging of spongy mesophyll cells of beech (*Fagus grandifolia*) leaves. **(A-C)** Representative live leaf areas dissected for isolation of spongy mesophyll cells of (A) control, (B) symptomatic beech leaf disease leaves with green, and (C) yellow interveinal patterns. **(D-F)** Spongy mesophyll cell shape of control (D) and symptomatic (E-F) leaf areas with different interveinal color patterns. White squares in the photomicrographs are magnified in D’-F’. Note the larger cell size for symptomatic leaves represented in red. **(G)** Average number of chloroplasts quantified in spongy mesophyll cells dissected from control (n = 25) and BLD symptomatic (n = 25) leaves. Due to high cell disruption and nematode activity, the chloroplasts of the mesophyll cell dissected from yellow interveinal areas were not quantified (F). **(H-J)** Representative individual spongy mesophyll cells of control (H), and green (I) and yellow (J) interveinal banding pattern of symptomatic leaves. Scale bars: 10 µm.

Interestingly, both dark green and yellowish BLD symptomatic areas presented ticker mesophyll layers in comparison to control beech leaves, accompanied by larger spongy mesophyll cells (see next section). These larger cells were also characterized by an increase of the chloroplast content (Fig 11G) when compared to control spongy mesophyll cells (Fig 11H-J). This overall increase of cell layers, and cell sizes, associated with an amplified number of chloroplasts, might explain the green interveinal patterns associated with symptomatic BLD leaves. While in green dark banding areas the presence of nematodes was limited, or totally absent in some cases, in acute symptomatic areas a large number of nematodes could be found (Fig 11F). In this last case, a large proportion of the spongy mesophyll cells presented disrupted cytoplasm and an apparent reduction/destruction of the chloroplasts (Fig 11J), probably influenced by the dynamic feeding activity of the nematodes from these cells. In the case of acute yellow-like interveinal bands, often related to the maturation of the nematode feeding site (Kanzaki et al. 2018), the disruption of cells by nematode feeding could explain their change of coloration.

### 8. Symptomatic interveinal leaf areas revealed an increased number of cell layers and augmented cell sizes

Finally, to track the overall structural changes occurring in mature symptomatic BLD leaves, we focused our following analyses in control (Fig 12A) and early autumn symptomatic BLD leaves (Fig 12D). Control mature beech leaves were flanked by single epidermal cell layers of both adaxial and adaxial sides of the leaf (Fig 12B-12C). The adaxial epidermis consisted of cells with relatively wavy anticlinal walls of variable size, whereas the abaxial layer was composed with similar cells but slightly smaller. The mesophyll consisted of one row of column palisade mesophyll cells on the adaxial side, and three layers of irregular shaped spongy mesophyll cells separated by intercellular air spaces towards the abaxial side of the leaf (Fig 12C). The adaxial and abaxial epidermis each occupied 14.7±2.3% to 11.08±1.72% of the total leaf thickness, respectively. The mesophyll tissue encompassed approximately 72.15% of total leaf thickness, of this the palisade is represented by 28.29±2.26%, while the spongy mesophyll is represented by 43.78±4.56%. Within the vasculature, a layer of compactly arranged sheath cells surrounds the water-bearing xylem cells adaxial and the sugar-bearing phloem cells abaxial (Fig 12C).

**Fig 12.**
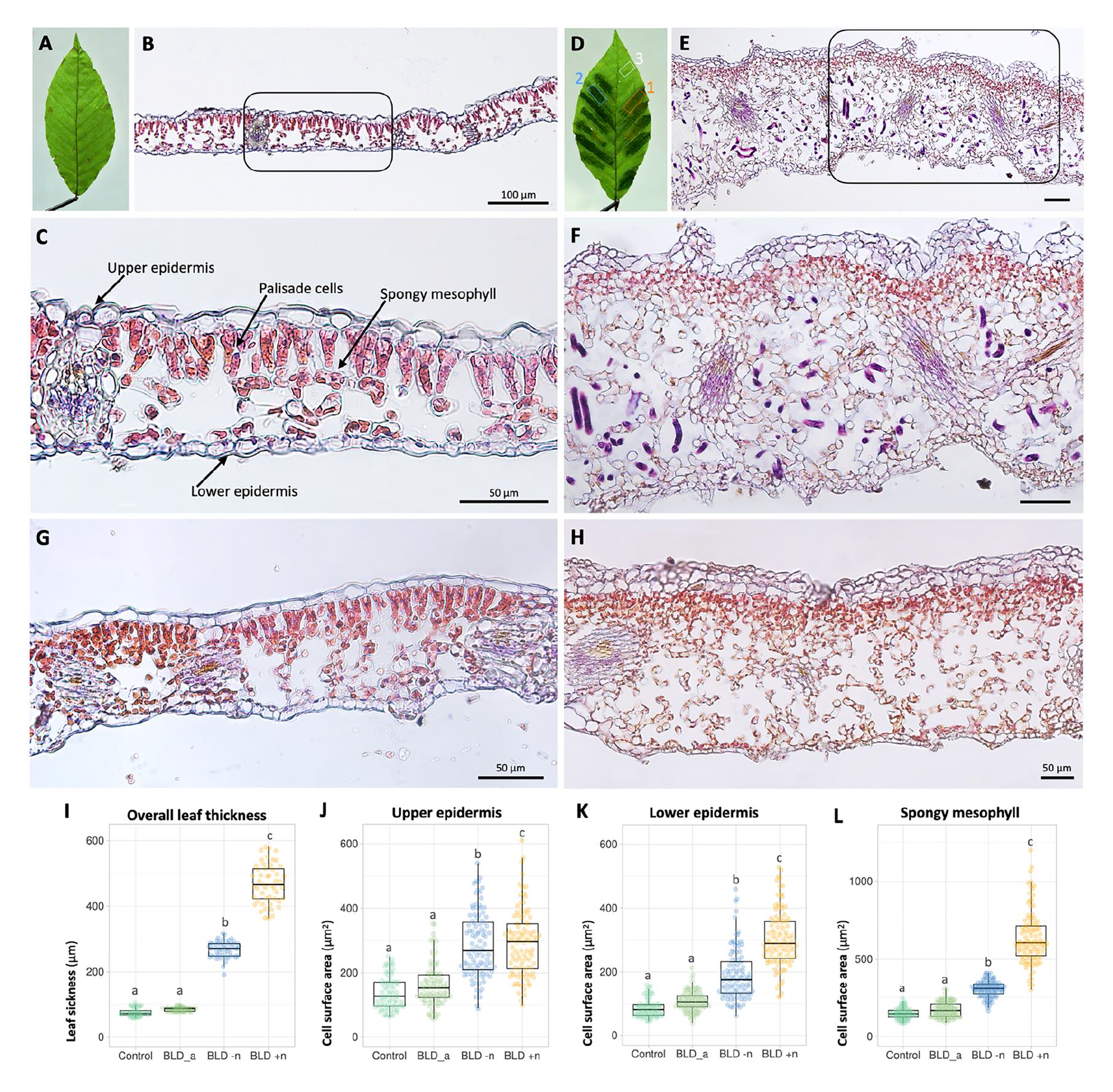
Cell architecture of beech (*Fagus grandifolia*) leaves. Photomicrographs of transverse sections of control and symptomatic beech leaf disease (BLD) leaves collected in early autumn. Sections were stained with a mixture of hematoxylin and eosin. **(A)** Representative control beech leaf. **(B-C)** Section of a mature control beech leaf showing differentiated cell layers as indicated in figure C. **(D)** Representative symptomatic BLD leaf. Dotted white, blue and orange squares represent sections of (G) asymptomatic, (H) green and (E-F) yellow banded areas of the same leaf used for histological analyses. **(E-F)** Sections of yellow symptomatic BLD areas showing extensive leaf thickness as a consequence of cell proliferation and cell hypertrophy. Nematode sections are stained (dark purple). **(G)** Section of a non-symptomatic area of a BLD leaf, which presents cell architecture similar to that of control leaves (B). **(H)** Section of a green interveinal area of a BLD leaf without nematodes. Note the additional cell layers in comparison to the control leaves (C). **(I)** Leaf thickness was measured from photomicrographs of control, asymptomatic (BLD_a) and symptomatic BLD areas without (BLD −n) and with (BLD +n) nematodes. For each condition a minimum of 50 measurements were made. **(J-L)** Cell surface (µm^2^) was measured for the upper epidermis (J), lower epidermis (K) and spongy mesophyll cells (L). Measures were made for control (n = 80 cells), asymptomatic (BLD_a, n = 100) and symptomatic BLD areas without (BLD −n, n = 100) and with (BLD +n, n = 100) nematodes. Box plots represents means ± SEM. Different letters indicate statistically significant differences compared with the control (*P* < 0.05). Scale bars: 100 µm (A, B); 50 µm (C-H).

From the same symptomatic BLD leaf, different areas were considered for histology, including: 1) yellowish/acute interveinal banding areas; 2) green/moderate interveinal banding areas, and 3) non-symptomatic areas, as highlighted in Fig 12D. Our analysis showed dramatic structural changes in symptomatic leaves (Fig 12E-12F). The acute areas of these leaves exhibited extensive proliferation of cells and cellular hypertrophy (Fig 12F) when compared to control leaves (Fig 12C). Both adaxial and abaxial epidermis vary from one to four layers of irregular shaped cells within the same leaf. The typical column palisade cells observed in uninfected leaves, converted into multiple layers of rounded cells packed with chloroplasts. A noteworthy difference was observed in the spongy mesophyll cellular composition, with a prominent and larger network of irregular-shaped cells and larger intercellular spaces, comprising up to 71.66±2.14% of the total thickness of the leaf (Fig 13F). The adaxial and abaxial epidermis each occupied 8.13±2.59% and 9.05±2.49% of the total leaf thickness, respectively, while the palisade cells represented by 8.98±1.56%. As a result of the significant increase of the number of cell layers, the overall thickness of these symptomatic areas increased approximately 500% in comparison to their normal size. In addition to the bulk of changes shown in the epidermis and mesophyll layers, the vascular bundles also presented apparently larger and a more complex number of cells in comparison to control. Associated with all these changes was the presence of numerous nematode sections within the intercellular spaces of the spongy mesophyll (Fig 12F), confirming our preceding analyses, and previously observations (Carta et al. 2023). Nematodes were not found between the palisade mesophyll cells nor in the vascular tissue of the leaf. Thus suggesting, that the spongy mesophyll cells are the main source of nutrients once nematode established within the leaves.

**Fig 13.**
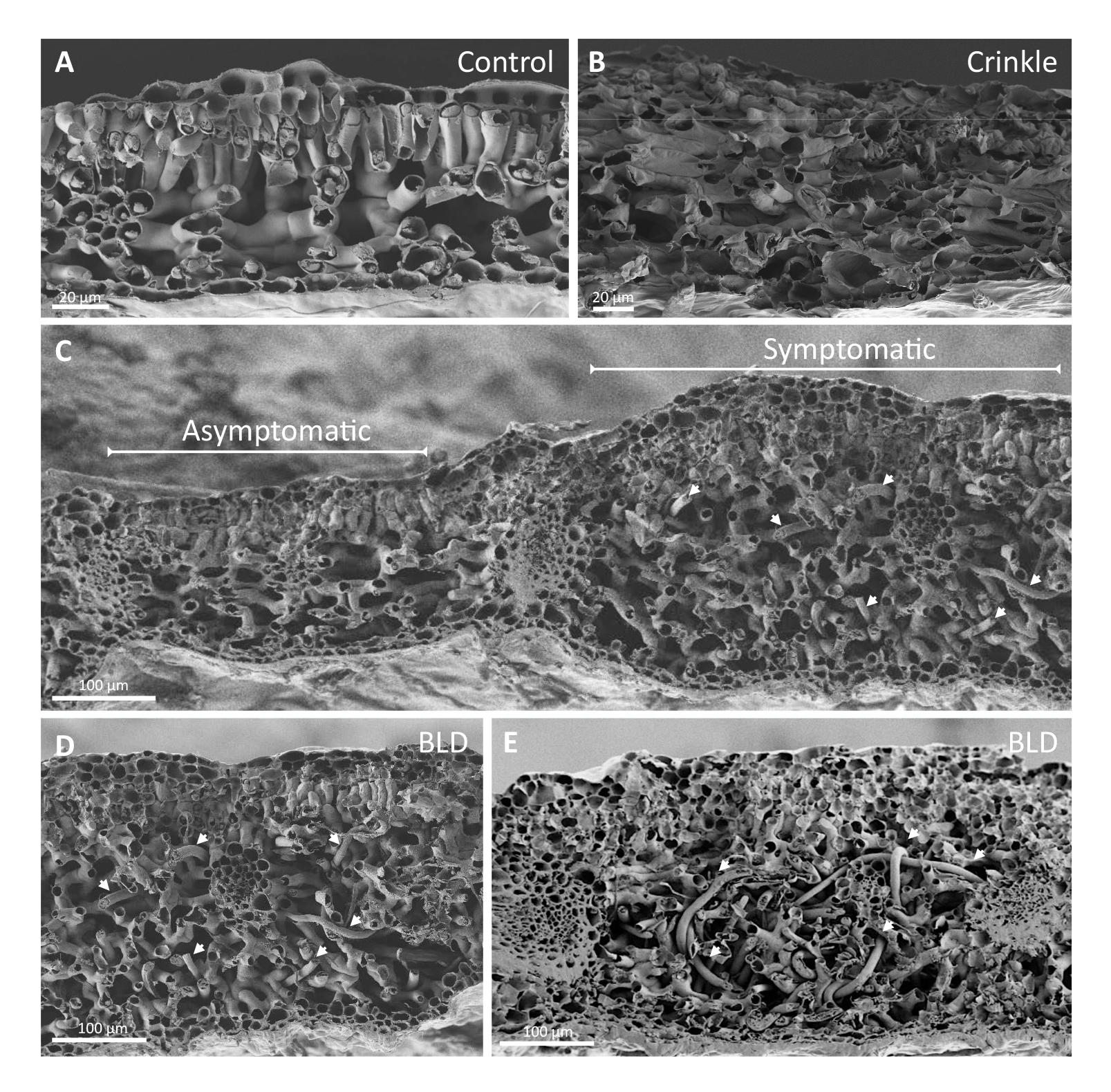
Scanning electron photomicrographs of transverse sections of control and symptomatic beech (*Fagus grandifolia*) leaves. **(A)** Transverse section of a control leaf showing the typical composition of six layers of cells. **(B)** Transverse section of a crinkled leaf (early spring) showing an abnormal number of cell layers and very compact intercellular spaces in the spongy mesophyll. **(C)** Representative transverse section of a symptomatic leaf showing asymptomatic (left) and BLD symptomatic (right) areas. **(D-E)** Symptomatic BLD section with numerous nematodes (arrows) within the spongy mesophyll. Scale bars: 20 µm (A, B); 100 µm (C-E).

Subsequently, we explored the anatomy of asymptomatic interveinal areas of surrounding BLD symptomatic areas within the same leaf. In these asymptomatic areas (Fig 12G) the overall leaf structure and thickness was compared with control leaves (Fig 12C). Interestingly, some dark green BLD banding areas of the examined leaves were devoid of nematodes. The cell organization, however, was very similarly to the nematode-infected interveinal leaf areas, i.e., comprising similar number of additional layers of cells (Fig 12H). The thickness of nematode-infected areas (468.68±59.72 µm) was significantly higher than those symptomatic areas lacking nematodes (268.62±25 µm), and to the thickness of both non-symptomatic (74.93±12.22 µm) and control (84.51±6.38 µm) leaves (Fig 12I).

We also measured individual cell sizes of epidermal (Fig 12J-12K) and spongy mesophyll (Fig 12L) cells of both uninfected and BLD asymptomatic and symptomatic leaf areas. Overall, the most dramatic changes were observed for the spongy mesophyll cells. On average mesophyll cell size of nematode infected areas (638.05±183.7 µm) was about twofold larger than those without nematodes (305.35±93.43 µm), and approximately four times larger than normal leaf cells (Fig 12L). Statistical analysis revealed that cells composing the other different layers of nematode-infected leaves were also significantly larger (*p* < 0.05) than those observed for control leaves with similar sizes. Based on our cell measurements, the spongy mesophyll cell enlargement suggests that the developmental program governing cell expansion are more severely affected by the presence of nematodes, which explains the difference of size to BLD symptomatic areas devoid of nematodes.

To validate our observations, SEM images were also used to visualize the cell layer structure of control, crinkle leaves collected in the spring, and non-symptomatic and symptomatic areas within the same BLD leaf collected in early autumn (Fig 13). Similarly, SEM revealed and highlighted the main cellular changes as described above, i.e., symptomatic BLD leaves were composed of irregular number of epidermal cell layers, followed by a dense distribution of circular palisade cells, and a large and complex network of spongy mesophyll cells embedding large number of nematodes, with more than the six cell layers observed in naïve leaves (Fig 13C-E). In contrast, while crinkle leaves displayed an increased number of cell layers, they presented very reduce inter-cellular spaces between the spongy mesophyll cells (Fig 13B) when compared to both control (Fig 13A) and other moderate BLD symptomatic leaves (Fig 13D-E). In summary, the formation of BLD symptomatic leaves is orchestrated by distinct cellular processes, involving an overall hyperplasia of the different cell layers of the leaf followed by augmented cell expansion, in particularly of the spongy mesophyll cells associated with the presence of the nematodes.

## Discussion

Although the origin of Lcm is still unresolved, the fast dispersal of this nematode can be associated with an apparent lack of resistance in native established forest trees suggests that this nematode was recently introduced to North America. Following this scenario, Lcm has successfully adapted to new environmental conditions and is thriving in this apparently new association with the American beech tree phenology. Understanding the complete life cycle of the nematode is necessary for comprehending its biology and interaction with its new, putative host species in their native range. This information is also critical for the implementation of adequate protective measurements for future monitoring, mitigation, and control of this disease. By characterizing the population dynamics of nematodes within the buds, we were able to provide additional data regarding the life cycle of this species (Kanzaki et al. 2018; Carta et al. 2020).

Our time-course observations demonstrated that the life cycle of Lcm is intimately associated with the developmental stages of the buds and leaves. The migration of nematodes into new developing buds is a critical step in their life cycle, not only as a source of nutrients, but also as a protective, overwintering structure. Similarly, other Anguinidae species use closed structures of the plant (e.g., seeds) for protection from hostile environmental conditions (Eisenback et al. 2013) or show highly adaptably to survive harsh environments, including cold tolerance (Svilponis et al. 2017; Carta et al. 2023). During the early interaction with the buds, Lcm actively feed on, and reproduce within the bud scales, suggesting that these tissues are sufficiently nourishing for nematode development; however, as the population increases, they interact with the shoot apex and leaf primordial cells which markedly alters leaf morphology.

Closed beech buds are highly frost-tolerant (Kreyling et al. 2015); however, the transition from active growth to dormancy is a part of the bud’s adaption to overwintering (Ruttink et al. 2007). This transition coincided with a generalized decline of nematode numbers during the colder months. With the onset of favorable conditions towards the spring, beech buds swell and break in early May, and by the middle of June, the leaves, formed in the bud in the previous year, fully expand and mature (Dengler et al. 1975). The low number of nematodes recovered from newly emerged leaves in the springtime resulted from the decrease of nematodes that survived the winter. This low number of nematodes associated with BLD leaves in the early months of the spring has been reported in other geographic areas in North America including those with much colder temperatures than in Virginia (Carta et al. 2020; Reed et al. 2020).

On the other hand, closed, poorly developed buds during early spring, contained a variable number of nematodes. Most of buds were surprisingly loaded with many nematodes and a large number of eggs, similar to that observed for other anguinids (Subbotin and Riley, 2012). Furthermore, a small fraction of buds revealed a significant number of dead nematodes with a low or total absence of eggs. This association of nematode numbers with different levels of symptoms suggest that the population levels above a certain threshold affect the development and ultimately the maturation of the leaf. In addition, our observations suggest that nematodes can reproduce and proliferate in large number within these buds. It is worth noting, that this type of information is important to consider when beech samples are processed for detection of nematodes, especially in the early spring, when the number of associated nematodes with symptomatic BLD leaves are still very limited. More importantly, the developing nematodes within these buds may serve as inoculum extending leaf infection.

The migration of Lcm from buds into newly emerged leaves seems to be a part of the natural development in the acquisition of nutrients in the loss of available bud tissues. As the leaves mature, nematodes are able to interact with a much larger volume of available plant tissue allowing for their exponential proliferation, exemplified by the hundreds of nematodes found in individual symptomatic leaves by summer’s end. This dynamic increase of nematodes will therefore contribute for the infection of new developing buds and perpetuating the spread of this disease in the following season.

Our characterization via microscopy showed that many of the cellular changes occurring in the bud scales resulted from cell hypertrophy rather by cell division, and that this transition involved the enlargement of the nuclei of these cells. These enlarged cells suggests that there is a potential link between the activation of the endoreduplication cycle with the establishment of these larger cells. The endoreduplication cycle is a modified cell cycle causing repeated DNA replication without nuclear division concomitant with cell expansion (Inzé and de Veylder, 2006; Kalve et al. 2014). The formation of enlarged cells are common features induced by many other plant-pathogens (Rodiuc et al. 2014; Favery et al. 2020) which stimulate a high metabolic activity to provide the parasite with suitable nutrient to complete their life cycle. Furthermore, the apparent absence of additional cell divisions in these tissues could correlate to the status of these cells no longer able to divide.

Although nematode-infected bud scales presented abnormal morphological changes, BLD is thought to be mainly a leaf disease. Since leaves are the primary organs for photosynthesis, they play an essential role for plant growth and development (Kalve et al. 2014). During primary morphogenesis, leaf development is sustained by successive oriented and coordinated cell division from the shoot apex throughout the formation of the leaf blade (Efroni et al. 2010). In *F. grandifolia*, 3 to 7 leaves are initiated on the shoot apex during July to August, afterwards, they are arrested in their development by late September resulting in characteristic winter buds (Dengler et al. 1975); however, depending on the geographical location and abiotic conditions, these periods can vary slightly. The early interaction of nematodes with the shoot apex meristem and leaf primordia occurs at this stage of extensive cell division within the closed bud and extends for long periods of time until the leaves breakthrough (i.e., from summer to the following spring). Our study showed that the interaction of nematodes in the earlier stages of leaf development was concomitant with an unexpected abnormal and extensive cell proliferation, resulting in a significant increase of the number of cell layers of the leaf. Targeting host cellular mechanisms through the modulation of the host cell cycle machinery is a recurrent strategy adapted by different plant-pathogens to induce specialized feeding sites (de Almeida Engler et al. 2015; Favery et al. 2020). Cell hyperplasia is a constant feature among the feeding sites induced by other Anguinidae (Palomares-Rius et al. 2017). Thus, it is likely that Lcm increases the rate or prolongs the time over which cell division occurs in the leaf.

Bud dormancy represents a survival strategy to persist during prolonged, unfavorable periods for growth. Dormancy results from the gradual transition from meristematic growth to a period of inactivity (Ruttink et al. 2007). Following this scenario, whether less active nematodes can induce cell division during bud dormancy still needs to be determined; however, as buds alter dormancy due to the return of favorable abiotic conditions, nematodes also resume their development. At this stage, young, non-infected leaves (<10 mm long) displayed the characteristic six, well-organized cell layers of not fully differentiated cells within the bud, as previously reported for *F. grandiflora* (Dengler et al. 1975). However, in nematode-infected buds, this normal leaf architecture was altered with additional and variable number of cell layers. The unique interveinal leaf symptoms suggest that these abnormal cell divisions were not synchronized along the full extent of the leaf, but rather from distinct rates of cell division that were stimulated by the inherent load of nematodes. The occurrence of randomly distributed, symptomatic interveinal bands that are present when the leaves break through the buds (Fearer et al. 2022) reinforces the idea that major changes in the morphology of the leaf must have occurred while the leaves were still within the bud. Support for this idea, when fully developed leaves were inoculated with nematodes, they failed to induce BLD symptoms, suggesting that the interaction of the nematodes within the bud plays a crucial step for the induction of BLD (Carta et al. 2020). Similarly, other Anguinidae species infecting aerial plant parts, including leaves, stems or inflorescences, induce their feeding site preferentially in actively growing undifferentiated tissues (Subbotin and Riley, 2012; Palomares-Rius et al. 2017). We speculate that abnormal cell proliferation results not only during the nematode interaction with the early leaf primordial cells, but also during the incessant interaction of Lcm with the leaf development within the bud. Overall, this ectopic cellular division of the developing leaves is a key mechanism to attain the variable upsurge of cells in symptomatic BLD leaves. However, the precise mechanism(s) leading to this abnormal ectopic cell division warrants further research.

Subsequently, during secondary morphogenesis of the leaf, cells stop dividing and begin to expand, resulting in additional growth of the leaves (Dengler et al. 1975; Kalve et al. 2014). Thus, correct regulation of cell expansion and proliferation fix the final size of the leaf (Vercruysse et al. 2020). Concomitant with the leaf maturation, the migration of nematodes, first onto the surface and then into the inner tissues of the leaf, results in the establishment of populations within the spongy mesophyll cell layers that increase substantially during the last part of summer and the first part of fall. The observation of an abnormal number of differentiated cell layers in mature BLD leaves provide further evidence of previous ectopic feeding stimulated mitotic activity in these interveinal leaf areas. The consequent hypertrophy of the spongy mesophyll cells associated with the presence of nematodes is probably a sign of endoreduplication occurring in these cells, similar to the cellular changes observed in the cells of the bud scales. For example, gall formation stimulated by the root-knot nematodes relies on polyploid cell formation where the nuclei of the giant cells are increased in number and in size from interference in the endocycle machinery (Vieira and de Almeida Engler et at. 2017).

One of the most striking features of BLD is the fine regulation of the cell machinery of the infected plant cells. Unlike sedentary cyst or root-knot nematodes, which interact directly with a relatively small number of root cells, Anguinidae species induce subtle changes, but in a much larger number of cells. This leads one to preclude that the changes in a larger number of cells relates to the nematode migratory status. In both cases, these specialized feeding sites act as a strong resource for plant metabolites and are often associated with extensive changes in plant growth and impose a considerable drain on the host performance (Rodiuc et al. 2014; Palomares-Rius et al. 2017). The deliberate induction of cell division and cell expansion of the host tissues suggests that these nematodes orchestrate the host cell cycle machinery. Evolutionary shifts between host plant lineages and Anguinidae species have been previously proposed, suggesting apparent evolutionary trend of gall evolution within this family (Subottin et al. 2004). Nevertheless, the overall cell composition of symptomatic BLD leaves is comparable, to a certain extent, to the cellular changes induced by other Anguinidae species, strongly supporting the link between the presence of Lcm and BLD. These changes consisted in extensive cell division and hypertrophy of the infected tissues (Palomares-Rius et al. 2017), which seems to be a fundamental step for the ontogeny and development of these nematode feeding sites. Because of the variety of the response by the plant, the processes and mechanisms are likely to be numerous. Surprisingly, and in contrast to root-knot and cyst nematodes (Favery et al. 2020; Goverse and Mitchum, 2022), the molecular mechanisms behind the formation of these anguinid nematode feeding sites are yet to be determined. In the same manner, the diversity of nematode effectors responsible for this (direct or indirect) fine tuning of the host cell manipulation within the Anguinidae is totally unknown.

Many unresolved questions about the fundamental processes of this new beech leaf disease remain. Further research at the molecular and biochemical levels are necessary to fully understand the context of the host responses to Lcm, as well as the potential interaction with other microorganisms. Furthermore, identification of nematode effector proteins and their potential host interaction may provide significant insight into the molecular pathways involved in the etiology of this very unique disease.

## Material and Methods

### Plant material collection

The area selected for our study is situated in Prince William Forest Park (Triangle, Virginia, US). This forest is characterized by a large proportion of beech trees and since the detection of BLD in 2021 (Kantor et al. 2022a), an increased number of symptomatic trees has been observed within this area. Leaves and buds were collected from *F. grandifolia* trees from October 2021 to April 2023. At each harvest time, several dozens of buds and/or leaves were collected from a minimum of ten randomly symptomatic BLD trees. Leaves with representative stages of BLD symptoms (i.e., slight, moderate, or acute interveinal dark green patterns) were collected at each harvest time. As control, buds and/or leaves were collected from similar asymptomatic size trees within the same forest park. Due to the high variability in leaf size across the growing seasons, leaves of comparable stages were collected from both control and symptomatic trees.

### Nematode extraction and quantification

Beech buds collected from control and BLD symptomatic trees were measured and weighed at different times of the year. Using sterile forceps, buds were dissected by removing individual bud scales and inner leaves in 1 ml of water in a 35 x 10 mm petri dish with a MVX10 Olympus stereoscope. Bud scales and leaves were rinsed, followed by three hours of incubation in distilled water at room temperature. Afterwards, nematodes were recovered and diluted in 2 ml of water. To quantify the total number of nematodes associated with each individual bud, three aliquots of 50 µl were placed in a concave microscope slide and nematodes were counted. Whole, individual scales or leaves of nematode-infected and control buds were photographed with a DP80 digital Olympus camera mounted on a MVX10 Olympus stereoscope. For the extraction of nematodes from mature leaves, BLD symptomatic leaves were selected and sectioned in 5 x 0.5 cm strips and placed in a 12 cm diameter Petri dish and submerged in water overnight. Afterwards, the solution containing the nematodes was poured into a 50 ml Falcon tube and centrifuged for 4 min at 3,000 g. The supernatant was removed, and the nematodes were used for morphological analyses. For morphological characterization nematodes were mounted in water agar slides (Eisenback, 2012) or fixed in 3% formaldehyde and processed to glycerin by the formalin glycerin method (Hooper, 1970; Golden, 1990). Fresh or fixed nematodes were photographed with a DP73 digital Olympus camera mounted on an Bx51 Olympus microscope.

### Detection of nematodes in beech twigs and DNA extraction

To verify the presence of nematodes on beech twigs, a total of 15 twigs of approximately 20-30 cm were randomly collected from five symptomatic BLD beech trees in early spring. All buds and leaves were extracted from the twigs, and each twig was then cut into 5 cm pieces long. The bark of the twigs was gently scratch with a scalpel in distilled water in a 12 cm diameter petri dish. Extracted nematodes were hand-picked and processed for morphological analyses or DNA extraction. Sets of five nematodes were placed in a 15 µl drops of digestion buffer (Invitrogen) in concave microscope slide and cut with a scalpel in several pieces. Nematodes sections were then transferred to a 1.5 ml tube. DNA extraction was performed with the PureLink Genomic DNA Kit (Invitrogen) in accordance with the manufacturer’s recommendations. Lcm detection was conducted with a Bio-Rad CFX OPUS 96 real-time PCR thermocycler targeting a fatty acid-and retinol-binding gene (FAR) using the primers 5’- GAAGCCCAAAGGAGATGAGAAG-3’ and 5’-TCCTCGGACAGAGCCTTATAC-3’. DNA extracted from a pool of nematodes collected from buds was used as positive control, while water was used as negative control.

### Histological sectioning and bright field microscopy

Freshly collected beech buds and leaves were considered for histological analyses using two different approaches. In the first approach, whole buds were fixed using a solution of 2% paraformaldehyde, 2.5% glutaraldehyde, 0.05% Tween-20, 0.05M sodium cacodylate, and 0.005M calcium chloride (pH=7). Buds were then post-fixed with 1% osmium tetroxide. The fixed buds were embedded in LX-112 resin (Ladd Research Industries) using a standard series of ethanol dehydration followed by infiltration with propylene oxide and then resin. All series and mixtures were graded at 25%, 50%, 75% and 100%, with the 100% steps performed in triplicate. All dehydration and infiltration steps were performed under vacuum in a microwave oven. Polymerization took place at 65°C for 48 hours. Sections were made using a Leica UC7 ultramicrotome equipped with a Histo Diamond Knife (DiATOME). Histological sections of 1 µm thickness were mounted onto a glass slide, dried at 65°C, and stained for 10 seconds with EMS-brand “epoxy tissue stain” (Electron Microscopy Sciences), which is a mixture of toluidine blue and basic fuchsin.

In a second approach, buds were processed after an initial assessment for the presence of nematodes by dissecting the first 4-5 scales. Afterwards, control and nematode-infected buds were fixed in 16% formaldehyde solution overnight at 4°C, and then dehydrated in an ethanol series (70%, 85%, 2 x 95% and 2 x 100%), followed by two washes with xylene, and embedded in glycol methacrylate resin Leica Historesin (Leica) following the protocol developed by Sampias and Rolls (https://www.leicabiosystems.com/knowledge-pathway/he-staining-overview-a-guide-to-best-practices/; Leica Biosystems; last accessed 3.10.2023). Paraffin embedded tissues were then sectioned with a thickness of 10 µm using a Shandon Finesse ME (Thermo Scientific) microtome. Slides were stained with hematoxylin and eosin staining using the Leica Autostainer ST5010 XL and covered slipped.

To study the cellular arrangements of symptomatic BLD leaves, symptomatic and non-symptomatic areas were dissected from the same individual leaves. Non-symptomatic leaves with similar sizes were processed as control. For histological analysis, control and BLD symptomatic leaves were initially fixed in 16% formaldehyde solution. Leaves were then embedded in glycol methacrylate resin Leica Historesin (Leica) and sectioned as described above (i.e., second approach). Sections of buds and leaves were photographed with a DP73 digital Olympus camera mounted on a Bx51 Olympus microscope. Outline drawings of leaf sections were post-processed using the Adobe Photoshop v. 2023 software. Cell and nuclei measurements were performed directly on bud or leaf sections using the CellSens ver. 1.6 software (Olympus). Data were plotted with PlotsOfData (Postma and Goedhart, 2019).

### Leaf tissue clearing and analyses of epidermal pavement cells

Analyses of the epidermal pavement cells of control and BLD symptomatic leaves was examined by processing leaves of similar sizes. Leaf clearing was performed by placing fresh leaves in 70% ethanol in a 50 ml tube at 75°C for 4 h, followed by a rehydration through an ethanol series (50%, 30%, 15%), and in a 25% (v/v) aqueous glycerol solution. The cleared tissues were then mounted in a glass slide and viewed under a Bx51 Olympus microscope equipped with a DP73 digital camera. Outline drawings of leaf epidermal pavement cells were post-processed using the Adobe Photoshop v. 2023 software.

### Fresh leaf spongy mesophyll cell measurements

For measuring the spongy mesophyll cell surface, fresh BLD symptomatic and control leaves were gently stripped with a scalpel in 20 ml of water in 12 cm diameter petri dish under a MVX10 Olympus stereoscope. Leaf sections were transferred to a glass slide and photographed with a DP73 camera attached to a Bx51 Olympus microscope. Cell measurements and chloroplast counting were performed using software CellSens ver. 1.6 (Olympus).

### Acid fuchsin staining of control and symptomatic BLD leaves

To follow up Lcm migration along symptomatic BLD leaves in the spring, fresh leaves were initially placed in 20 ml of acid fuchsin solution [20 ml of distilled water, 500 µl of acid fuchsin (0.15 g/10 ml of water) and 500 µl of glacial acetic acid] (Bybd et al. 1983) and briefly warmed in a microwave, followed by 2 h at room temperature. Leaves were afterwards gently washed with distilled water, and nematodes associated with the leaves were quantified under a MVX10 Olympus stereoscope. To stain nematodes within BLD symptomatic leaves early in the fall, comprising both symptomatic and non-symptomatic areas, leaf strips of approximately 5 x 0.5 cm were excised and distained in 70% ethanol at 75°C overnight. Leaf fragments were then placed in 20 ml of acid fuchsin solution in a 50 ml plastic tube and placed under a CentriVap Cold Trap vacuum (Labconco) at 4°C for 1 h. Leaf fragments were then rinsed three times in distilled water and distained in a clearing solution (equal volumes of lactic acid, glycerol, and distilled water) overnight at room temperature. Leaves were afterwards mounted in a slide for imaging in a 25% (v/v) aqueous glycerol solution. Similar preparations were performed for control leaves.

### Scanning electron microscopy of beech leaves

Preparation of fresh leaf samples for scanning electron microscopy (SEM) was performed as previously described (Hall and Hawes, 1991). Briefly, leaves were fixed overnight at 4°C in 40 ml of fixative (2.8% glutaraldehyde in 0.1M sodium cacodylate and 0.02% triton X-100), followed by 3 x 15 min washes with buffer (0.1M sodium cacodylate). Sample dehydration was performed through an ethanol series (25%, 50%, 70%, 85%, 95%, 3 x 100%) for 15 min each. Specimens were kept overnight at 4°C in a covered jar with a lid. Samples were then transferred to CPD carriers and placed in a glass specimen jar filled with fresh 100% ethanol. Critical point drying was performed using a Leica EM CPD300 instrument (Leica). Leaf samples were mounted to aluminum stubs with conductive tabs (carbon tape) and coated by adding a drop of colloidal silver to the edge of each sample and then sputter coated with 10nm Au/Pd (20 Angstrom thickness). Imagens were captured with a Zeiss Sigma Field Emission Scanning Electron Microscope (Zeiss). Healthy and BLD diseased leaves were also prepared for low temperature-SEM (LT-SEM) imaging using a Hitachi SU7000 Schottky-Emitter SEM (Hitachi High Technologies America). The SEM was outfitted with a Quorum PP2000 cryo-prep chamber (Quorum Technologies). Fresh leaves were divided into 1 mm^2^ specimens using surgical scissors. Leaf specimens were mounted on a copper plate in two rows, with either the upper or lower leaf surface exposed. Samples were mounted in a thin layer of cryo-adhesive applied to the plate, into which samples were gently pressed. The copper plate was then immersed in liquid nitrogen to fix the adhesive and freeze the sample. To maintain a clean, visible surface, the samples were then exposed to a partial vacuum in the slushing chamber to solidify the liquid nitrogen. The plate was then transferred (*in vacuo*) onto a −140°C cryo-stage inside of a Quorum preparatory chamber. Following *in vacuo* sublimation, the entire plate was sputter coated with platinum inside the preparatory chamber. The sample was then transferred into the main body of the LT-SEM onto the central cryo-stage, which was maintained at a constant temperature of −140°C. Images were taken using a 5kV accelerating voltage and low probe current, with 256 seconds for each integration.

### Statistical analyses

All data were analyzed using analysis of variance (ANOVA), and means were compared using Tukey’s honestly significant difference (HSD) test at the 5% probability level. In all figures, statistical significance (*P* < 0.05) is indicated as different letters.

## Acknowledgments

We thank Margaret McDonald and Joseph Mowery (USDA, Beltsville, MD) for technical support. The co-authors would also like to acknowledge the Huck Institutes’ Microscopy Core Facility for use of the Zeiss Sigma Field Emission Scanning Electron Microscope and John Cantolina for helpful discussions on sample preparation (Penn State University, University Park, PA). This study was supported by the United States Department of Agriculture Agricultural Research Service CRIS project number 8042-22000-322-000D (PV, ZAH). This research was partially funded by Penn State University through the Huck Institutes of the Life Sciences (MRK).

## Author contributions

**Conceptualization:** Paulo Vieira

**Formal analysis:** Paulo Vieira, Mihail R. Kantor, Andrew Jansen, Zafar A. Handoo, Jonathan D. Eisenback

**Methodology:** Paulo Vieira, Mihail R. Kantor, Andrew Jansen, Zafar A. Handoo, Jonathan D. Eisenback

**Project administration:** Paulo Vieira

**Writing – original draft:** Paulo Vieira, Jonathan D. Eisenback

**Writing – review and editing:** All co-authors

## Notes

### Competing Interest Statement

The authors have declared no competing interest.

